# Inhibitor-2 directs formation of PP1 holoenzymes through a docking motif-dependent transfer of catalytic subunits to adapters

**DOI:** 10.64898/2026.03.15.711929

**Authors:** Neha Varshney, Aleesa J. Schlientz, Johnathan L. Meaders, Karen Oegema, Arshad Desai

## Abstract

The catalytic subunits of protein phosphatase 1 (PP1) achieve spatiotemporal substrate specificity by assembling with diverse regulatory adapters to form holoenzymes. Three conserved proteins—Sds22, Inhibitor-2 and Inhibitor-3—facilitate loading of PP1 catalytic subunits (PP1cs) onto adapters. We show here that Inhibitor-2 is central to a dynamic cycle that directs formation of adapter-bound PP1 holoenzymes. Inhibitor-2 engages PP1cs via two adapter-like docking motifs (RVxF and SILK) and an active site-binding inhibitory region. While Inhibitor-2 depletion produced moderate phenotypes, mutation of its RVxF docking motif caused severe defects resembling global PP1c inhibition. The RVxF mutant did not prevent PP1c binding or reduce PP1c stability but inhibited formation of adapter-bound holoenzymes. The severe effects of the RVxF mutation were suppressed by linked mutation of the inhibitory active site-binding motif. These results suggest that Inhibitor-2 is integral to a dynamic cycle that delivers PP1cs to adapters, with its RVxF motif being critical for coupling relief of active site inhibition to adapter handoff.

## INTRODUCTION

Protein Phosphatase 1 (PP1) is a ubiquitous eukaryotic enzyme responsible for nearly half of all serine/threonine dephosphorylation events (Ceulemans and Bollen, 2004; Cohen, 2002). In contrast to the large protein kinase family, PP1 activity is encoded by a small number of catalytic subunits (PP1cs). The functional versatility of PP1 arises from the association of these catalytic subunits with regulatory adapters, also referred to as RIPPOs (Regulatory Interactors of Protein Phosphatase One; (Cao et al., 2022; Lemaire and Bollen, 2020)), to form an array of PP1 holoenzymes that act in diverse biological contexts (Bollen et al., 2010; Virshup and Shenolikar, 2009). Adapters bind PP1cs by employing short (4–6 residue) linear sequence motifs, most prominently the RVxF motif, that are typically located in unstructured regions that dock onto conserved PP1c surfaces (Hendrickx et al., 2009; Heroes et al., 2013; Nilsson, 2019; Peti et al., 2013). Adapters target PP1cs to distinct subcellular locations and contribute to substrate specificity; they also possess domains and interactions encoding functions unrelated to PP1c docking. The sharing of a limited set of catalytic subunits across a diverse array of adapters highlights the highly dynamic combinatorial control of PP1 (Bollen, 2001).

Prior to their association with adapters to form diverse holoenzymes, PP1 catalytic subunits associate with a set of highly conserved proteins—Sds22, Inhibitor-2 (Inh2), and Inhibitor-3 (Inh3)—that play critical roles in the biogenesis of active subunits and their handoff to adapters (Cao et al., 2022; Lemaire and Bollen, 2020; Verbinnen et al., 2017). Nascent PP1 first associates with Sds22 and Inh3, whose coordinated action promotes the formation of active, metal-loaded PP1cs (Choy et al., 2019; van den Boom et al., 2023; Weith et al., 2018). Inh2, by contrast, has long been characterized as a PP1 inhibitor; this inhibition is explained by direct active site occlusion by a conserved segment of Inh2 (Cohen, 1989; Huang and Glinsmann, 1976; Hurley et al., 2007; Lemaire and Bollen, 2020). Early biochemical studies further suggested that Inh2 can act as a chaperone promoting PP1c activation, with phosphorylation of a conserved threonine playing a role (Alessi et al., 1993; Ballou et al., 1983; Hemmings et al., 1982; Park and DePaoli-Roach, 1994). Genetic studies have implicated Inh2 function in multiple processes involving phosphorylation-dephosphorylation cycles (Eto et al., 2002; Lemaire et al., 2024; Peel et al., 2017; Tung et al., 1995; Wang et al., 2008a; Wang et al., 2008b) and support Inh2 promoting PP1c functions *in vivo*. One recent study in human cells proposed that Inh2 functions as a stabilizer for specific PP1c-adapter holoenzymes, and its absence selectively affects the phosphorylation status of the substrates of these holoenzymes (Lemaire et al., 2024).

Inh2 binds PP1cs using multiple conserved motifs, including SILK and RVxF docking motifs that overlap with adapter-binding interfaces in holoenzymes, as well as an active site-binding inhibitory sequence (HYNE) embedded within the longer IDoHA (Inhibitor-2 <u>Do</u>cking site for the <u>H</u>ydrophobic and <u>A</u>cidic grooves) motif (Hurley et al., 2007; Lemaire and Bollen, 2020). The fact that both Inh2 and many PP1 adapters utilize SILK and RVxF class motifs to dock onto the PP1c surface suggests a competitive handoff mechanism with formation of transient trimeric complexes (Dancheck et al., 2011; Eto et al., 2002). However, there is limited understanding of how the insights derived from biochemical analysis relate to Inh2 functions *in vivo* and whether Inh2 serves as a global buffer of PP1 activity, as a selective facilitator of the assembly of specific holoenzymes, or as an active participant in a dynamic cycle that facilitates the loading of PP1c onto a diverse set of adapters to form functional holoenzymes.

In this study, we use the *C. elegans* early embryo to explore the functional relationship between PP1c and its conserved regulators, with a focus on Inh2. The *C. elegans* genome encodes 4 PP1c isoforms, two of which, PP1cα^GSP-2^ and PP1cβ^GSP-1^, are broadly expressed; the other two isoforms are selectively expressed during spermatogenesis (Wu et al., 2012). As anticipated from PP1’s major role in cellular processes, PP1cα^GSP-2^ and PP1cβ^GSP-1^ contribute to diverse processes in early embryos, including meiotic chromosome segregation, nuclear re-assembly, control of mitotic duration, centriole duplication and embryo polarization (Calvi et al., 2022; Hattersley et al., 2016; Hsu et al., 2000; Kim et al., 2017; Nadarajan et al., 2021; Peel et al., 2017; Tzur et al., 2012). Conserved PP1 regulatory proteins have also been implicated in early embryonic processes, for example, Sds22 in polarity establishment (Li et al., 2025) and Inh2 in centriole duplication (Peel et al., 2017).

Here, we show that PP1c catalytic subunits are largely redundant in the one-cell *C. elegans* embryo and, consistent with current thinking, Sds22 and Inh3 are important for PP1c biogenesis and activity. In contrast to Sds22 and Inh3, Inh2 makes a more selective contribution to PP1c functions, with its depletion resulting in modest phenotypes. Analysis of double depletions suggests that Inh2 makes a greater functional contribution to PP1cβ^GSP-1^ than PP1cα^GSP-2^. Unexpectedly, specific mutation of Inh2’s RVxF docking motif resulted in a severe defect resembling global loss of PP1c functions; by contrast, mutation of the SILK and active site-binding HYNE motifs had little effect. This result supports a central role for Inh2 in a dynamic cycle that facilitates the loading of PP1cs onto a diverse set of cellular adapters, with the Inh2 RVxF motif acting as a critical element that couples relief of PP1c catalytic inhibition to adapter handoff.

## RESULTS

### Functional analysis of PP1 catalytic subunits and their core regulators in the one-cell *C. elegans* embryo

The two embryonically expressed PP1c isoforms, PP1cα^GSP-2^ and PP1cβ^GSP-1^, exhibit high sequence and structural similarity, differing primarily in their C-terminal tails (**Fig. S1A**). To assess their individual functions, we designed dsRNAs that enabled selective RNAi-mediated depletion of each PP1c, as assessed by imaging of embryos expressing *in situ* GFP-tagged PP1cα^GSP-2^ and PP1cβ^GSP-1^ (**Fig. S1B-D**). While individual depletions of PP1cα^GSP-2^ or PP1cβ^GSP-1^ did not significantly perturb the first embryonic division, depletion of both catalytic subunits led to catastrophic meiotic failure and subsequent mitotic defects (**Fig. 1A, Movie S1**). These results indicate that the two PP1c subunits are largely redundant for PP1 functions in the one-cell *C. elegans* embryo.

**Figure 1.**
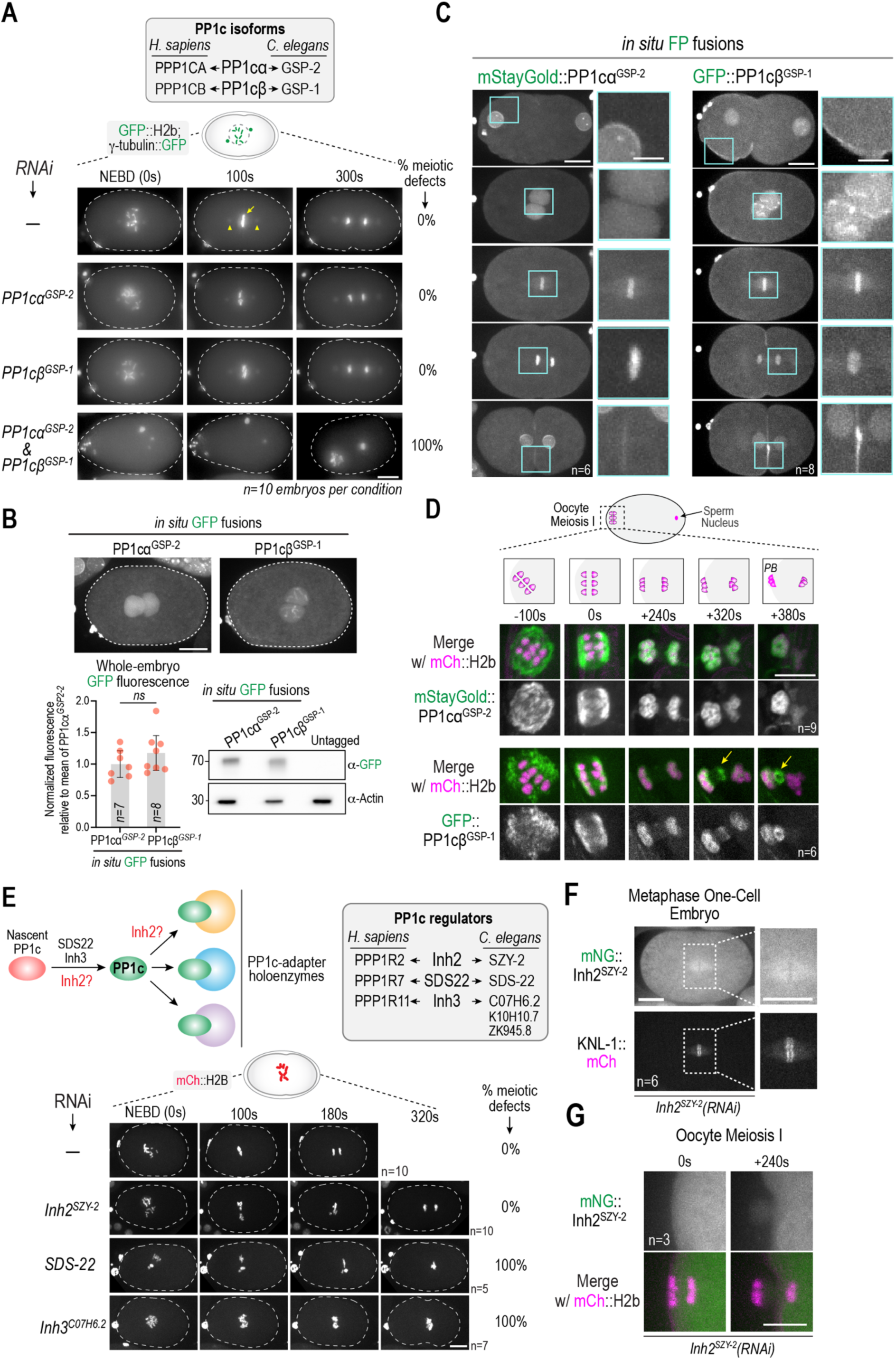
A functional survey of PP1 catalytic subunits and conserved regulators in the one-cell *C. elegans* embryo. **(A)** (*top*) PP1 catalytic subunit genes in humans and *C. elegans*. See also *Fig. S1A*. (*bottom*) Stills from timelapse movies of one-cell embryos expressing fluorescently-tagged histone H2b and γ-tubulin, which label chromosomes (*arrow*) and spindle poles (*arrowheads*), respectively, for the indicated conditions (see also *Fig. S1B-D*). Meiotic failure was scored based on presence of excess chromatin in one-cell embryos and improperly formed polar bodies. *n* is the number of embryos imaged per condition; Scale bar, 10 µm. **(B)** Quantification of expression of the two PP1c isoforms. (*top*) Example images of one-cell embryos expressing *in situ* GFP-tagged fusions imaged under identical conditions. (*bottom left*) Quantification of whole embryo fluorescence. N2 (*wildtype*) untagged embryos were imaged in parallel to correct for embryo autofluorescence (*see Fig. S1B*). (*bottom right*) Immunoblot of indicated worm strain extracts. Actin serves as a loading control. Error bars are the 95% confidence interval; p-value (ns=not significant) is from a Mann-Whitney test. Scale bar, 10 µm. **(C)** Localization of the two PP1cs at different stages of the first embryonic division. Cyan boxes highlight regions magnified to the right. Scale bar, 10 µm for whole embryo images; 5 µm for magnified region. **(D)** Localization of the two PP1cs on the oocyte meiosis I spindle, with mCh::H2b-labeled chromosomes. Schematics above depict chromosome distribution at the timepoints shown. The mSG::PP1cα^GSP-2^ strain also expressed an mCherry-tagged plasma membrane marker. Scale bar, 5 µm. **(E)** (*top*) Schematic highlighting the three conserved PP1c regulators: Sds22, Inhibitor-2 (Inh2) and Inhibitor-3 (Inh3)–and their encoding genes in humans and *C. elegans*. (*bottom*) Stills from timelapse movies of one-cell embryos with fluorescently-labeled chromosomes for the indicated conditions. See also *Fig. S1E,F*. *n* is the number of embryos imaged per condition; meiotic defects were scored as in (*A*); Scale bar, 5 µm. **(F) & (G)** Images of mNeonGreen (mNG)-tagged Inh2^SZY-2^ in mitotic metaphase (*F*) and oocyte meiosis I anaphase (*G*). For mitotic metaphase imaging, the strain expressed an mCherry fusion with the kinetochore marker KNL-1. For meiotic imaging, the strain expressed an mCherry fusion with histone H2b and an mCherry-tagged plasma membrane marker. *n* is number of embryos imaged. Scale bars, 10 µm for whole embryo images and magnified region in (*F*) and 5 µm for (*G*).

We assessed the relative amounts of the two PP1c isoforms by quantifying the fluorescence signal of *in situ* GFP-tagged PP1cα^GSP-2^ and PP1cβ^GSP-1^ in one-cell embryos (**Fig. 1B; Fig. S1B**). The quantification revealed that PP1cα^GSP-2^ and PP1cβ^GSP-1^ are expressed at near-equivalent levels, a conclusion supported by immunoblotting (**Fig. 1B**). We did not observe substantial compensation in expression of PP1cα^GSP-2^ following depletion of PP1cβ^GSP-1^ or vice versa (**Fig. S1C,D**). Monitoring the localization of the two catalytic subunits by imaging *in situ* tagged mStayGold::PP1cα^GSP-2^ and GFP::PP1cβ^GSP-1^ fusions revealed shared and specific subcellular localizations for the two isoforms in the early embryo (**Fig. 1C**). Both PP1cs localized to the nucleoplasm, kinetochores, and anaphase chromatin. PP1cα^GSP-2^ was additionally observed on the nuclear envelope and at nucleoli, whereas PP1cβ^GSP-1^ localized prominently to the embryo cortex, prophase mitotic chromatin, and the midbody (**Fig. 1C, Movie S2**). The two PP1cs also localized to the spindle area and on anaphase chromosomes during oocyte meiosis I (**Fig. 1D; Movie S3**). PP1cβ^GSP-1^ was additionally detected at the late midbody ring during extrusion of the first polar body at the end of meiosis I (*yellow arrows*; **Fig. 1D, Movie S3**). Thus, the two PP1c catalytic subunits are expressed at near-equivalent levels and exhibit shared and specific localization patterns in the one-cell embryo and during oocyte meiosis. Depletion of either individual subunit reduces the total amount of PP1c by about 50% and is largely tolerated during the early embryonic divisions.

Next, we imaged embryos following depletion of the *C. elegans* orthologs of the conserved regulators important for PP1c biogenesis and function: Sds22 (SDS-22 in *C. elegans*; (Li et al., 2025; Peel et al., 2017), Inh2 (SZY-2 in *C. elegans* (Peel et al., 2017)) and Inh3. There are three genes encoding Inh3-related proteins in *C. elegans*, only one of which (*C07H6.2*) is expressed in early embryos (Hashimshony et al., 2015). Consistent with prior work establishing a central role for SDS-22 and Inh3 in PP1c biogenesis (Choy et al., 2019; Weith et al., 2018), depletion of SDS-22 or Inh3^C07H6.2^ resembled double depletion of the catalytic subunits (**Fig. 1E**). Monitoring PP1cβ^GSP-1^ additionally revealed a significant reduction in PP1c levels in SDS-22 depleted embryos (**Fig. S1E**). By contrast, Inh2^SZY-2^ depletion had a significantly milder effect, with the primary phenotype being modestly delayed anaphase onset (**Fig. 1E**) and slowed chromosome decondensation.

As efforts to tag Inh2^SZY-2^ at its endogenous locus were unsuccessful, we generated a single-copy transgene insertion composed of the endogenous Inh2^SZY-2^-encoding locus fused to mNeon Green (mNG). mNG::Inh2^SZY-2^ localized diffusely to the cytoplasm and concentrated in the nucleus (**Fig. S1F)**. Following RNAi targeting the endogenous locus, mNG::Inh2^SZY-2^ fluorescence was no longer detected, indicating efficient depletion (**Fig. S1F**). To enable molecular replacement experiments, we generated a second mNG::Inh2^SZY-2^-encoding transgene that was re-encoded at the nucleotide level to render it RNAi-resistant (**Fig. S1G**; *see also Fig. S4A*). The RNAi-resistant transgene-encoded mNG::Inh2^SZY-2^ rescued the partial embryonic lethality observed following endogenous Inh2^SZY-2^ depletion (**Fig. S1G**), indicating that the fusion is functional. Following endogenous Inh2^SZY-2^ depletion, mNG::Inh2^SZY-2^ was not detected at prominent sites of PP1c localization in one-cell embryos or during oocyte meiosis I (**Fig. 1F,G**), indicating that Inh2^SZY-2^ is not a stable component of PP1 holoenzyme complexes.

Overall, a functional survey in one-cell *C. elegans* embryos revealed redundancy of the equally expressed PP1c α and β isoforms, confirmed central roles for SDS-22 and Inh3 in PP1c biogenesis, and suggested a more specific contribution of Inh2^SZY-2^, which does not concentrate at sites of PP1c action. We thus decided to focus on understanding how Inh2^SZY-2^ acts to support PP1c functions.

### Inhibitor-2 contributes to the function of both PP1c isoforms but makes a greater contribution to PP1cβ

To understand the role of Inhibitor-2 and its relationship to the PP1c α and β isoforms, we capitalized on events during the first embryonic division of the *C. elegans* embryo to investigate the Inh2^SZY-2^ depletion phenotype and the effect of Inh2^SZY-2^ removal on PP1cα and PP1cβ. In embryos depleted of Inh2^SZY-2^ or individually depleted of PP1cα^GSP-2^ or PP1cβ ^GSP-1^, no major defects were observed prior to nuclear envelope breakdown (NEBD). We therefore focused on measuring the interval between NEBD and anaphase onset, which is controlled by removal of an inhibitory phosphorylation on the anaphase activator CDC-20 (Kim et al., 2017). This dephosphorylation event involves PP1c bound to its kinetochore-localized adapter KNL-1, acting on CDC-20 as it fluxes through kinetochores (Kim et al., 2017). Individual depletions of PP1cα^GSP-2^ or PP1cβ^GSP-1^ led to a similar small increase in the NEBD to anaphase interval (from ∼180s to ∼220s) consistent with the KNL-1 adapter forming holoenzymes with both PP1c isoforms (Espeut et al., 2012). Depletion of Inh2^SZY-2^ had a more significant effect than depletion of PP1cα^GSP-2^ or PP1cβ^GSP-1^ alone (NEBD to anaphase ∼270s), suggesting that Inh2^SZY-2^ contributes at least partially to the function of both isoforms (**Fig. 2A**). The effects of depleting Inh2^SZY-2^ were similar to the effects of preventing the recruitment of PP1c to kinetochores through mutation of the KNL-1 adapter; both perturbations delayed anaphase onset and impaired chromosome alignment (**Fig. S2A-E; Movie S4** (Espeut et al., 2012; Kim et al., 2017)). As expected for a delay caused by reducing the rate of removal of an inhibitory phosphorylation on CDC-20, replacement of CDC-20 with a mutation that renders it non-phosphorylatable on its inhibitory site significantly reduced the anaphase onset delay (**Fig. S2F**).

**Figure 2.**
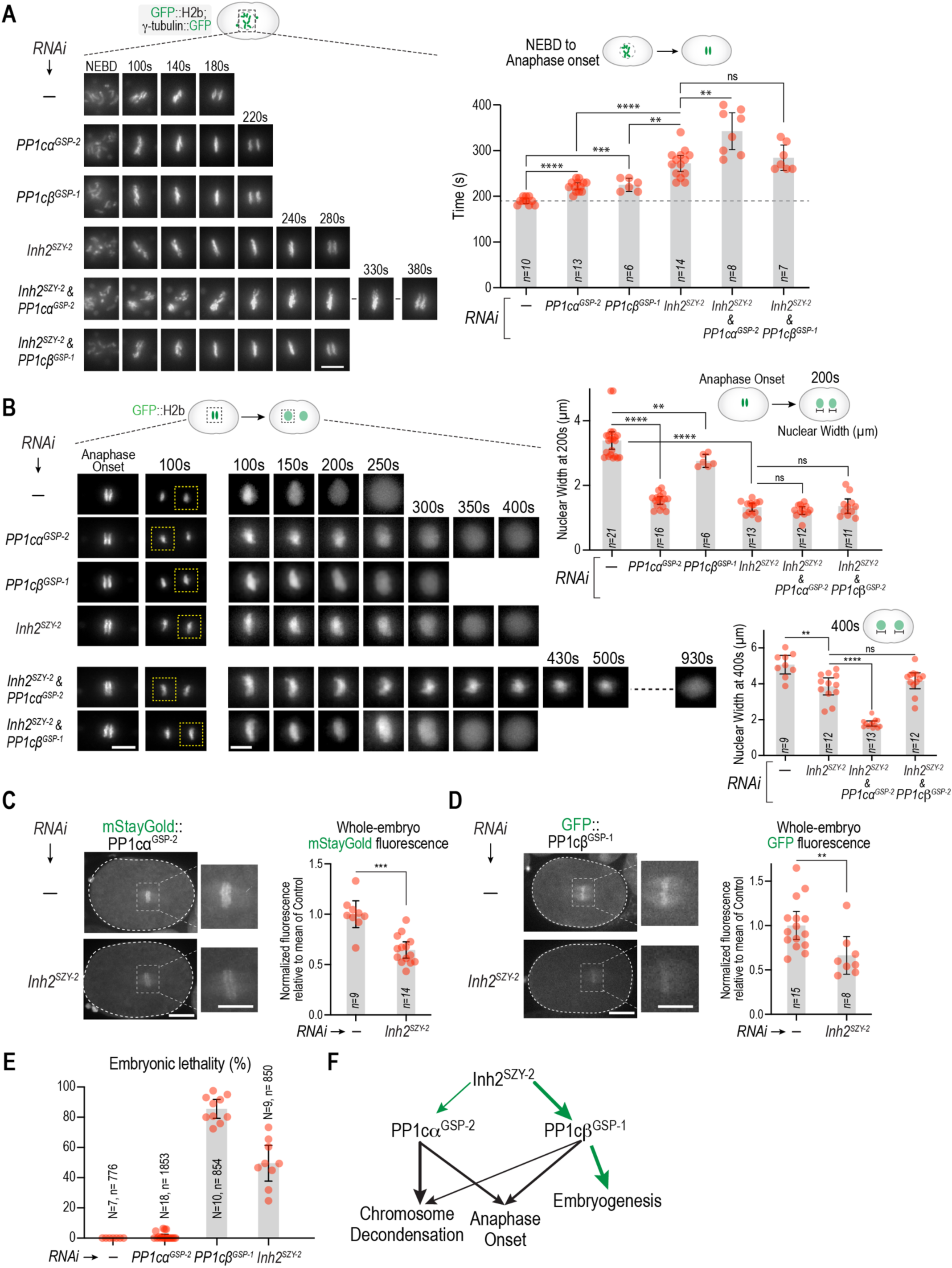
Inhibitor-2 contributes to the function of both PP1 catalytic subunits but has a greater impact on PP1cβ function. **(A)** (*left*) Stills from timelapse movies showing chromosome dynamics during mitosis for the indicated conditions. (*right*) Measurement of the NEBD to anaphase onset interval in one-cell embryos for the indicated conditions. Scale bar, 5 µm. See also *Fig. S2A-G*. **(B)** (*left*) Stills from timelapse movies showing chromosome decondensation after anaphase onset in one-cell embryos. One of the two reforming nuclei, indicated with the dashed box, is magnified to the right. Scale bars, 5 µm. (*top right*) Graph plotting nuclear width 200s after anaphase onset; (*bottom right*) Graph plotting nuclear width 400s after anaphase onset. **(C, D)** (*left*) Images and (*right*) quantification of *in situ*-tagged PP1cα^GSP-2^ (*C*) or PP1cβ^GSP-1^ (*D*) whole embryo fluorescence for the indicated conditions, measured as described in *Fig. S1B*. Scale bar, 10 µm for whole embryo images and 5 µm for magnified images. **(E)** Analysis of embryonic lethality for the indicated conditions. *N* is number of worms analyzed and *n* is the number of embryos scored. Lethality of embryos laid 24-48h after injection of dsRNA into late L4 stage larvae was scored. Note that the No RNAi and Inh2^SZY-2^ depletion data are the same as in *Fig. S1G.* See also *Fig. S2G*. In graphs in panels (*A*)-(*D*), *n* is the number of embryos imaged. All error bars are the 95% confidence interval. p-values are from Mann-Whitney tests (ns=not significant; **=p<0.01; ***=p<0.001; ****=p<0.0001). **(F)** Schematic summary of the functional contributions of the two PP1c isoforms in the indicated processes and of Inh2^SZY-2^ to the functions of each isoform.

To assess the relative contribution of Inh2^SZY-2^ to the function of the α and β isoforms, we combined Inh2^SZY-2^ depletion with depletion of individual PP1c subunits. Depletion of PP1cα^GSP-2^ along with Inh2^SZY-2^ led to a significant extension of the NEBD to anaphase onset interval compared to depletion of Inh2^SZY-2^ alone (∼340s vs. ∼270s), indicating that PP1cα^GSP-2^ retains substantial function in Inh2^SZY-2^-depleted embryos. By contrast, depletion of PP1cβ^GSP-1^ along with Inh2^SZY-2^ did not significantly increase the NEBD to anaphase onset interval, suggesting that PP1cβ^GSP-1^ function is compromised by the absence of Inh2^SZY-2^. Additional support for the idea that PP1cβ^GSP-1^ has a greater dependence on Inh2^SZY-2^ for function than PP1cα^GSP-2^ came from analysis of chromosome decondensation after anaphase onset (**Fig. 2B**), a second PP1-dependent process requiring reversal of mitotic phosphorylation (Antonin and Neumann, 2016) mediated by adapters such as RepoMan in vertebrates and, potentially, the nucleoporin MEL-28/ELYS in *C. elegans* (Hattersley et al., 2016; Qian et al., 2011; Trinkle-Mulcahy et al., 2006; Vagnarelli et al., 2006; Vagnarelli et al., 2011). Individual depletion of PP1cα^GSP-2^ slowed decondensation, quantified by measuring the width of the chromatin mass at a fixed time point (200s) after anaphase onset, significantly more than PP1cβ^GSP-1^ depletion (**Fig. 2B**), suggesting that the adapter that controls chromosome decondensation preferentially uses PP1cα^GSP-2^ over PP1cβ^GSP-1^, although PP1cβ^GSP-1^ can support decondensation at a slowed rate when PP1cα^GSP-2^ is depleted. Inh2^SZY-2^ depletion significantly delayed decondensation. Similar to the analysis of the NEBD to anaphase onset interval, co-depletion of PP1cα^GSP-2^ along with Inh2^SZY-2^ significantly slowed chromosome decondensation compared to Inh2^SZY-2^ depletion alone, indicating that PP1cα^GSP-2^ retains the ability to promote chromosome decondensation in Inh2^SZY-2^-depleted embryos (**Fig. 2B, Movie S5**). By contrast, co-depletion of PP1cβ^GSP-1^ and Inh2^SZY-2^ did not further slow decondensation compared to Inh2^SZY-2^ depletion alone, indicating that PP1cβ^GSP-1^ largely lacks the ability to perform this function in the absence of Inh2^SZY-2^ (**Fig. 2B, Movie S6**).

One potential explanation for the greater reliance of PP1cβ^GSP-1^ on Inh2^SZY-2^ compared to PP1cα^GSP-2^ is that Inh2^SZY-2^ loss could affect the stability or expression of PP1cβ^GSP-1^ more than that of PP1cα^GSP-2^. To test this idea, we quantified whole embryo fluorescence of the two *in situ* GFP-tagged PP1c isoforms following Inh2^SZY-2^ depletion (**Fig. 2C,D**). Both PP1c isoform levels were equivalently reduced (on average by ∼40%) following Inh2^SZY-2^ depletion (**Fig. 2C,D**), indicating that, while Inh2^SZY-2^ is required for wildtype PP1c expression levels, it does not selectively affect PP1cβ^GSP-1^ over PP1cα^GSP-2^.

The conclusion that PP1cβ^GSP-1^ function is more reliant on Inh2^SZY-2^ than PP1cα^GSP-2^ received additional support from analysis of embryonic lethality (**Fig. 2E**). RNAi-mediated depletion of PP1cα^GSP-2^ alone did not lead to significant embryonic lethality while depletion of PP1cβ^GSP-1^ or Inh2^SZY-2^ both caused significant lethality. These data suggest that PP1cβ^GSP-1^ plays an essential role in later stages of embryogenesis that cannot be compensated for by PP1α^GSP-2^. The fact that Inh2^SZY-2^ depletion also leads to significant embryonic lethality (**Fig. 2E; Fig. S2G**) is consistent with Inh2^SZY-2^ supporting PP1cβ^GSP-1^-specific functions in embryogenesis.

Collectively, the analysis above indicates that, while Inh2^SZY-2^ contributes to the function of both PP1c isoforms, PP1cα^GSP-2^ retains significant function when Inh2^SZY-2^ is depleted whereas PP1cβ^GSP-1^ is more severely compromised (**Fig. 2F**).

### Mutation of Inhibitor-2 motifs involved in PP1c binding reveals an unexpectedly severe phenotype

Structural analysis of the human Inh-2–PP1c complex has shown that Inh2 employs three motifs – SILK, RVxF and HYNE – to engage distinct surfaces of PP1c (Hurley et al., 2007). On one face of the PP1c surface, the SILK and RVxF motifs of Inh2 dock into conserved pockets that are also used by adapters—such as KNL-1, MEL-28 and PAR-2 in *C. elegans* (Calvi et al., 2022; Espeut et al., 2012; Hattersley et al., 2016)—during the formation of PP1c holoenzymes. Inh2 wraps around PP1c to its opposite side, where the HYNE motif—a conserved 4-amino acid sequence embedded within the longer IDoHA (Inhibitor-2 Docking site for Hydrophobic and Acidic grooves) region of Inh2—binds and inhibits the PP1c active site (Hurley et al., 2007; Lemaire and Bollen, 2020). Thus, any handoff of PP1c from an Inhibitor-2–PP1c complex to adapters like KNL-1 and MEL-28 will require the SILK and RVxF motifs of the adapters to displace the equivalent motifs on Inh2. This displacement must also be coordinated with dissociation of the Inh2 HYNE motif from the active site and the release of Inh2. To visualize the interfaces made by the three motifs of *C. elegans* Inh2^SZY-2^ with PP1cα^GSP-2^ and PP1cβ^GSP-1^, we employed AlphaFold 3 (**Fig. 3A; Fig. S3A-E**). The structural modeling predicted interfaces consistent with prior structural analysis of the human complex, providing a structural basis for the mutational analysis described below.

**Figure 3.**
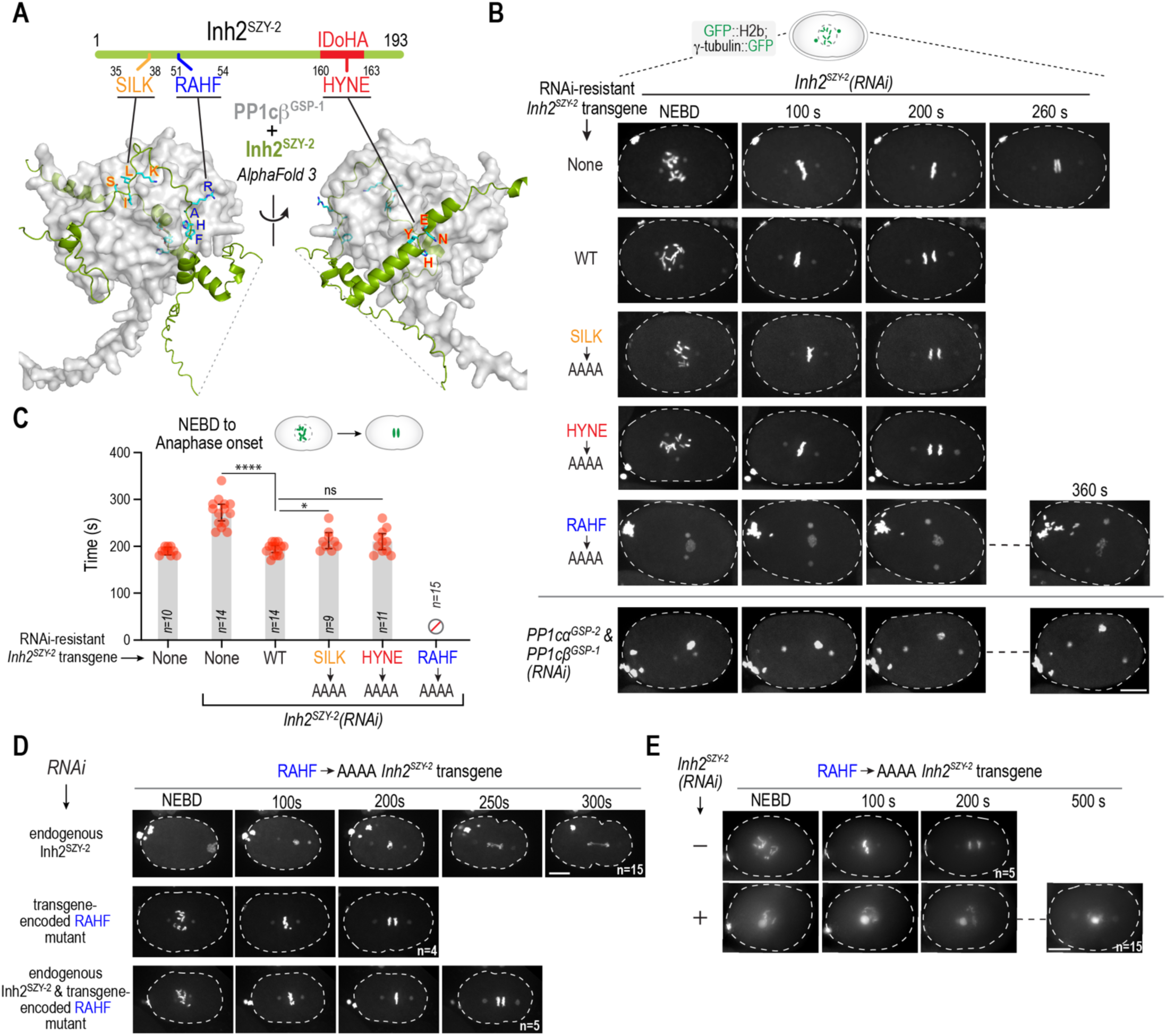
Mutation of the RVxF motif of Inhibitor-2 results in a severe phenotype resembling global loss of PP1 function. **(A)** Schematic of Inh2^SZY-2^ (*top*) and two views of an Alphafold 3 model of the Inh2^SZY-2^–PP1cβ^GSP-1^ complex (*bottom*), highlighting the three conserved PP1c-binding motifs: SILK, RVxF and HYNE; dashed line indicates an unstructured region predicted with low confidence. See also *Fig. S3*. **(B)** Stills from timelapse movies of one-cell embryo mitosis expressing fluorescently tagged histone H2b and γ-tubulin for the indicated conditions. See also *Fig. S4A-D*. Note that the PP1c double depletion embryo shown on the bottom is one of the 10 embryos from Fig. 1A. Scale bar, 5 µm. **(C)** Quantification of the NEBD-anaphase onset interval for the indicated conditions. *n* is the number of embryos filmed; error bars are the 95% confidence interval; p-values are from Mann-Whitney tests (ns=not significant; *=p<0.05; ****=p<0.0001). The values plotted for the first two conditions (no transgene, no RNAi; no transgene, *Inh2^SZY-2^(RNAi)*) are the same as in Fig. 2A. Note that for the RAHF◊AAAA mutant, the severity of the phenotype precluded measurement of the NEBD-anaphase onset interval. **(D)** Stills from timelapse movies of one-cell embryo mitosis expressing fluorescently tagged histone H2b and γ-tubulin for the indicated conditions. To deplete the transgene-encoded RAHF mutant form of Inh2^SYZ-2^, a dsRNA homologous to the recoded region of the transgene (*Fig. S4A*) was employed for *RNAi.* Note that the top embryo (endogenous Inh2^SZY-2^ depletion, transgene-encoded RAHF mutant) is one of the 15 RAHF mutant embryos from panel (*B*). Scale bar, 10 µm. See also *Fig. S4E*. **(E)** Stills from timelapse movies of one-cell embryo mitosis expressing fluorescently tagged histone H2b and γ-tubulin for the indicated conditions. Note that the bottom embryo (endogenous Inh2^SZY-2^ depletion, transgene-encoded RAHF>AAAA mutant) is one of the 15 RAHF mutant embryos from (*B*). Scale bar, 10 µm.

To gain insight into how Inhibitor-2 contributes to the formation of PP1 holoenzyme complexes, we used an Inh2^SZY-2^ RNAi-resistant transgene system to replace endogenous Inh2^SZY-2^ with mutants in each of the three interacting motifs (**Fig. S4A**). We first confirmed that the WT transgene is functional by showing that it rescues embryonic lethality and the anaphase onset delay caused by Inh2^SZY-2^ depletion (**Fig. 3B,C; Fig. S4B**). Individual mutation of the Inh2^SZY-2^ SILK and HYNE motifs had relatively little phenotypic impact: the NEBD-anaphase onset interval was largely unaltered (**Fig. 3B,C; Fig. S4B; Movie S7**) and chromosome decondensation, which we analyzed for the HYNE mutant, was normal (**Fig. S4C,D**). Unexpectedly, mutation of the Inh2^SZY-2^ RVxF motif led to severe defects that resembled the co-depletion of both PP1c isoforms (**Fig. 3B; Movie S7**). Given the degree of phenotypic severity, we were unable to quantitatively monitor the effect of the Inh2^SZY-2^ RVxF mutation on the NEBD-anaphase onset interval or chromosome decondensation. This surprising result indicated that mutation of the Inh2^SZY-2^ RVxF motif results in a significantly more severe phenotype, similar to loss of PP1c function, than depletion of Inh2^SZY-2^. To confirm that the unexpectedly severe phenotype observed for the Inh2^SZY-2^ RVxF mutant was indeed due to the transgene-encoded mutant, we employed two dsRNAs for RNAi: one targeting endogenous Inh2^SZY-2^ and the second targeting the transgene-encoded RVxF mutant (whose coding sequence was altered for RNAi-resistance). Removing the transgene-encoded mutant along with endogenous Inh2^SZY-2^ reverted the severe phenotype to the significantly milder Inh2^SZY-2^ depletion phenotype (**Fig. 3D; Fig. S4E**). We additionally assessed whether the severe phenotype was dominant or was only observed when endogenous Inh2^SZY-2^ was depleted and found that it required endogenous Inh2^SZY-2^ depletion (**Fig. 3E**). Thus, *in vivo* mutational analysis of the three conserved PP1c-interacting motifs of Inh2^SZY-2^ revealed an unexpectedly severe phenotype for the RVxF motif mutant of Inh2^SZY-2^ that resembled global loss of PP1c function.

### Linked mutation of both RVxF and the active-site binding HYNE motif suppresses the severe phenotype of RVxF-mutant Inhibitor-2

One potential explanation for the surprisingly severe phenotype of the RVxF mutant of Inh2^SZY-2^ is that this mutant form blocks a dynamic cycle of Inh2^SZY-2^ association and dissociation with PP1cs required for forming adapter-bound PP1c holoenzymes. In other words, when the RVxF interface is disrupted, the entire PP1c pool is irreversibly trapped in an Inh2^SZY-2^ RVxF mutant-bound inactive state. This model has two predictions: first, the RVxF mutant of Inh2^SZY-2^ should retain the ability to bind and inhibit PP1c employing the SILK and HYNE motifs; second, mutating the HYNE motif, which associates with the enzymatic active site to block activity, in the RVxF mutant would likely compromise PP1c binding and relieve active site inhibition, thereby reversing the severe phenotype of the RVxF mutant. To test these two predictions, we first employed yeast two-hybrid analysis to monitor binding of Inh2^SZY-2^ to PP1cα^GSP-2^ (**Fig. 4A**). The single motif mutants all interacted with PP1cα^GSP-2^, with the RVxF mutant exhibiting robust interaction and the HYNE mutant exhibiting a modestly compromised interaction (**Fig. 4A**). In contrast, the double RVxF-HYNE mutant was significantly compromised in binding (**Fig. 4A**). These binding results suggested that the RVxF-HYNE double mutant should behave similarly to Inh2^SZY-2^ depletion *in vivo*, in contrast to the significantly more severe RVxF mutant.

**Figure 4.**
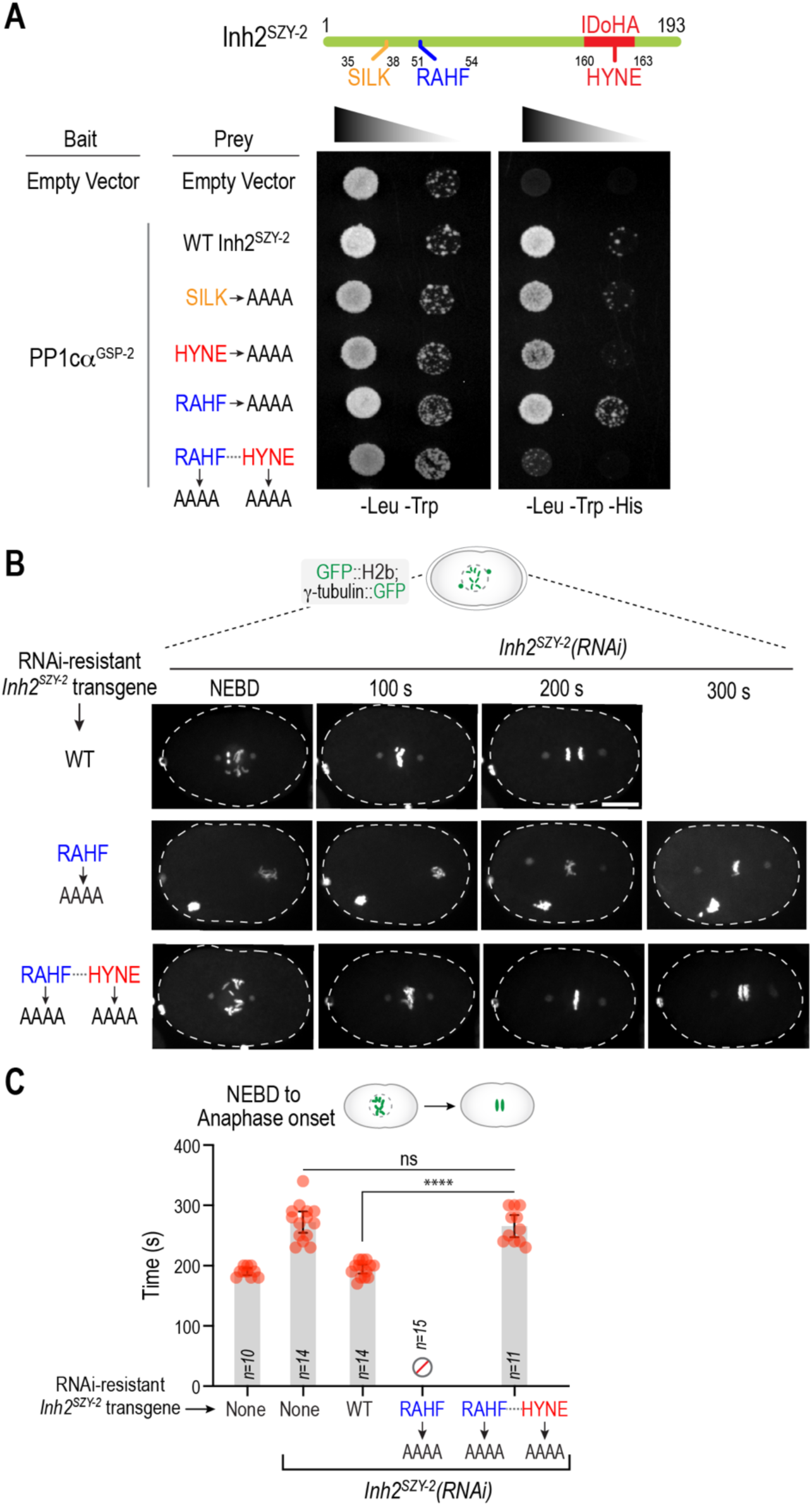
Linked mutation of the active-site binding motif suppresses the severe phenotype of the RVxF mutant of Inhibitor-2. **(A)** Yeast two-hybrid analysis of the interaction of WT and indicated motif mutant variants of Inh2^SZY-2^ with PP1cα^GSP-2^. The results shown are representative of 2 independent experiments. **(B)** Still from timelapse movies of mitosis for the indicated conditions. Note that the middle embryo (endogenous Inh2^SZY-2^ depletion, transgene-encoded RAHF mutant) is one of the 15 RAHF mutant embryos from Fig. 3B. Scale bar, 10 µm. **(C)** Quantification of the NEBD-anaphase onset interval for the indicated conditions. *n* is the number of embryos filmed; error bars are the 95% confidence interval; p-values are from Mann-Whitney tests (ns=not significant; *=p<0.05; ****=p<0.0001). The data plotted for the no transgene, WT transgene and RAHF mutant conditions are the same as in Fig. 2A and Fig. 3C. Note that for the RAHF◊AAAA mutant, the severity of the phenotype precluded measurement of the NEBD-anaphase onset interval.

To test this prediction, we generated a transgene insertion expressing the double RVxF-HYNE mutant and compared it to the single RVxF mutant. Consistent with the binding analysis, the double RVxF-HYNE mutant did not exhibit the severe phenotype observed for the RVxF mutant (**Fig. 4B; Movie S8**). Quantification of the NEBD-anaphase onset interval indicated that the double RVxF-HYNE mutant resembled depletion of Inh2^SZY-2^ (**Fig. 4C**). Thus, the severe defects caused by the Inh2^SZY-2^ RVxF mutant can be suppressed by linked mutation of the active site-binding HYNE motif.

### The Inhibitor-2 RVxF motif mutation prevents localization of PP1c to its sites of action

The diverse localization patterns of PP1cs reflect the formation of holoenzyme complexes with adapters that concentrate at specific cellular locations, such as the meiotic spindle and mitotic kinetochores (**Fig 1C, D**). The results above suggest that the RVxF mutant of Inh2^SZY-2^ prevents the formation of these holoenzyme complexes, potentially by trapping PP1cs in a bound, inactive state that cannot be transferred to adapters. An alternative model is that the RVxF mutant triggers degradation of PP1cs resulting in the observed severe phenotypes. To distinguish between these two models, we imaged *in situ* mStayGold-tagged PP1cα^GSP-2^ in the presence of WT, RVxF mutant or RVxF-HYNE double mutant Inh2^SZY-2^, following endogenous Inh2^SZY-2^ depletion. As the RVxF mutant causes severe meiotic defects that are compounded as the first mitotic division proceeds, we focused the analysis on oocyte meiosis I. We first quantified cytoplasmic fluorescence to determine if the RVxF mutant caused PP1c degradation. This analysis revealed that the RVxF mutant did not cause any reduction in PP1cα^GSP-2^ levels (**Fig. 5A**), while the RVxF-HYNE double mutant, which compromises binding to PP1cs, resembled Inh2^SZY-2^ depletion in lowering PP1c levels (**Fig. 5A**). Thus, the severe phenotype of RVxF mutant Inh2^SZY-2^ cannot be attributed to PP1c degradation.

**Figure 5.**
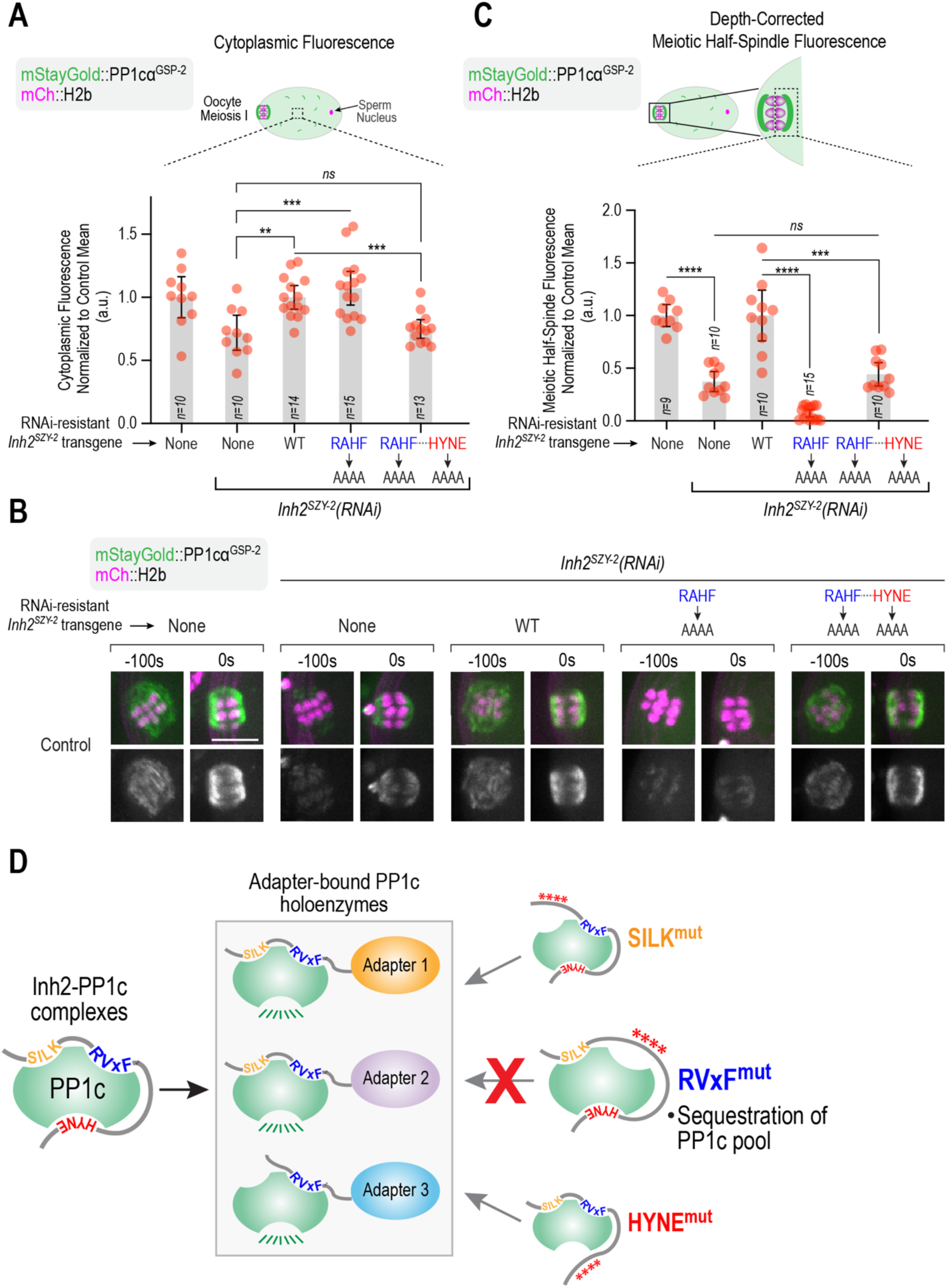
The RVxF mutant of Inhibitor-2 sequesters PP1 catalytic subunits preventing formation of adapter-bound holoenzymes. **(A)** Analysis of PP1cα^GSP-2^ levels for the indicated conditions, measured by quantification of cytoplasmic mStayGold-fused PP1cα^GSP-2^ fluorescence. The *Inh2^SZY-2^(RNAi)* measurements were normalized to the mean value of no RNAi controls; the RAHF and RAHF-HYNE mutant measurements were normalized relative to the WT transgene mean. *n* is number of embryos imaged; p-values are from Mann-Whitney tests (ns=not significant; **=p<0.01; ***=p<0.001). **(B)** Images of the meiotic spindle region for the indicated conditions. The strains imaged also expressed an mCherry-tagged plasma membrane marker. See also *Fig. S4F*. Scale bar 5 µm. **(C)** Quantification of meiotic half-spindle mStayGold::PP1cα^GSP-2^ fluorescence for the indicated conditions. The half-spindle on the oocyte cytoplasmic side was measured. The mStayGold fluorescence measurements were depth-corrected using the mCh::H2b signal. The depth-corrected *Inh2^SZY-2^(RNAi)* measurements were normalized relative to mean value of the No RNAi controls. The RAHF and RAHF-HYNE mutant transgene depth-corrected measurements were normalized relative to WT transgene controls. *n* is number of embryos imaged; p-values are from Mann-Whitney tests (ns=not significant; ***=p<0.001). **(D)** Schematic summary of the key findings. See text for details.

We next focused on the meiotic spindle, where PP1cs are known to concentrate (**Fig. 1D**; (Hattersley et al., 2016)). Depletion of Inh2^SZY-2^ reduced meiotic spindle-associated PP1cα^GSP-2^ by ∼50% and this reduction was rescued by expression of WT Inh2^SZY-2^ (**Fig. 5B,C**). Notably, the Inh2^SZY-2^ RVxF mutant caused a severe depletion of spindle-associated PP1cα^GSP-2^, while the double RVxF-HYNE mutant did not (**Fig. 5B,C**). Thus, the Inh2^SZY-2^ RVxF mutant binds to and sequesters PP1c, preventing it from forming meiotic spindle-localized adapter-bound holoenzymes. Consistent with this, a fluorescently tagged version of the Inh2^SZY-2^ RVxF mutant, did not ectopically localize to adapter-concentrated sites, such as the meiotic spindle (**Fig. S4F**).

Collectively, these results support a model in which Inh2^SZY-2^ dynamically cycles on and off PP1cs, a process that is blocked by the Inh2^SZY-2^ RVxF mutant, which sequesters the entire PP1c pool into an Inh2^SZY-2^ RVxF mutant-bound inactive state. These results also highlight a critical role for the RVxF motif of Inh2^SZY-2^ in the handoff of PP1cs from Inh2 to adapters and the concomitant release of inhibition that enables active holoenzyme formation.

## DISCUSSION

While Inh2 was discovered in the late 1970s as a heat-stable inhibitor of phosphorylase phosphatase (Huang and Glinsmann, 1976), its precise contributions to PP1 functions *in vivo* have proven challenging to elucidate. Inh2 is a potent inhibitor of PP1cs in purified systems and early biochemical experiments demonstrated a phosphorylation-based mechanism acting on Inh2-PP1c complexes to generate active PP1cs (Cohen, 1989; Lemaire and Bollen, 2020). A recent study in human cells has implicated Inh2 in the formation of specific PP1 holoenzymes (Lemaire et al., 2024). More generally, studies across experimental systems suggest that Inh2 behaves as a promoter of PP1 functions in the cellular context, in contrast to its initial biochemical identification as an inhibitor of PP1 activity.

The work we present here employing the *C. elegans* embryo contributes to efforts focused on defining the role of Inh2 in promoting PP1 functions. Our most striking result is that mutating the RVxF class docking motif of *C. elegans* Inh2 leads to the sequestration and global inhibition of PP1cs (**Fig. 5D**); by contrast, individual mutation of its two other PP1c docking motifs, SILK and active site-occluding HYNE, had little effect (**Fig. 5D**). RVxF mutant Inh2 did not cause PP1c degradation, distinguishing it from depletions of the PP1c biogenesis factors like Sds22 (Cao et al., 2024). The severe phenotype of the RVxF mutant could be reverted to the milder Inh2 loss-of-function phenotype by linked mutation of the active-site binding HYNE motif. Consistent with these *in vivo* results, the Inh2 RVxF mutant maintained the ability to interact with PP1cs, while the RVxF-HYNE double mutant did not; retention of PP1c-binding by Inh2 RVxF mutants has also been reported in other systems (Freville et al., 2013; Lemaire et al., 2024). Based on these results, we propose that Inh2 associates with PP1cs transiently as part of a dynamic cycle, in which the RVxF motif is critical for both adapter handoff and relief of active site inhibition (**Fig. 5D**). This idea is consistent with a recent proposal that adapters with an RVxF motif may be guided to Inh2-associated PP1cs by the Inh2 RVxF motif (Lemaire et al., 2024). How the phosphorylation cycle of Inh2 described in *in vitro* biochemical experiments (Cohen, 1989; Lemaire and Bollen, 2020) relates to the observations we present here remains to be clarified.

Our results also suggest that PP1c isoforms exhibit different degrees of dependence on Inh2 for productive association with adapters. While our data suggests that Inh2 promotes the association of both PP1c isoforms with adapters, in the absence of Inh2, PP1cα^GSP-2^ appears to make more productive associations with adapters than PP1cβ^GSP-1^. We note that PP1cα^GSP-2^, but not PP1cβ^GSP-1^, harbors a “TPPR” motif in its C-terminus that has been suggested to act as an intramolecular active site inhibitor (Lemaire et al., 2024); phosphorylation of the threonine residue in the TPPR motif is one of two mechanisms known to downregulate PP1 activity during mitosis (Dohadwala et al., 1994; Kwon et al., 1997; Moura et al., 2025). We speculate that, the presence of an intramolecular inhibitory region in the C-terminal tail may increase the extent to which a PP1c isoform can access an Inh2-independent pathway to adapter association and activation.

Overall, the work presented here suggests that Inh2 contributes to the formation of diverse PP1c holoenzymes. A major future challenge will be to address the precise mechanistic basis for why the RVxF mutant of Inh2 sequesters PP1cs and how that informs the normally coordinated cycle of active site inhibition release and adapter handoff to form functional holoenzymes.

## MATERIALS AND METHODS

### C. elegans *Strains*

*C. elegans* strains used in the study are listed in **Table S1**. All strains were maintained at 20°C on nematode growth media (NGM) plates seeded with OP50-1 *E. coli*. All transgenes including RNAi-resistant, tagged with mNG or untagged, were generated by single-copy insertions at specific chromosomal loci using the MosSCI method (Frokjaer-Jensen et al., 2008). Briefly, the *Inh2^SZY-2^*genomic locus – either endogenous or re-coded for RNAi-resistance, with or without a fluorescent protein tag, and under control of the *mex-5* promoter and *tbb-2* 3’ UTR – was cloned into the pCFJ151 vector. The integration plasmids were mixed with a plasmid expressing the transposase (pCFJ601) and co-injection marker plasmids pCFJ190, pCFJ104, pGH8, and pMA122 used to screen against extrachromosomal arrays; the plasmid mix was injected into both gonad arms of strain EG6699 to generate single-copy insertions in chromosome II (Frokjaer-Jensen et al., 2008). Transformants were selected based on their ability to rescue the mobility defect of the parental strains. Injected worms were singled onto NGM plates and grown for 7-10 days before being heat shocked for 3 hrs at 34°C. Surviving progeny were screened for the absence of extrachromosomal array markers (mCherry), and singled onto fresh plates. Successful integration was assessed by PCR, and integrated cassettes were sequence-validated using Oxford Nanopore longread sequencing.

For the Inh2^SZY-2^ RNAi replacement system, a region of the coding sequence was modified to introduce silent mutations to render them resistant to a dsRNA targeting the endogenous gene. A gene block containing the reencoded Inh2^SZY-2^ was purchased from Genewiz (from Azenta LifeSciences) and cloned into pCFJ151.

*In situ* GFP-tagged PP1cβ^GSP-1^ and PP1cα^GSP-2^ were previously described (Kim et al., 2017). Endogenous PP1cα^GSP-2^ was tagged with monomeric Staygold (mSG; (Hirano et al., 2022; Ivorra-Molla et al., 2024) at the N-terminus using CRISPR/Cas9 (Paix et al., 2015). A Cas9-RNP mix, containing the guide sequence for PP1cα^GSP-2^ (5’-GACGTAGAAAAGCTTAATC-3’), repair template, Cas9 (purchased from Berkeley MacroLab), and guide for the co-injection marker were injected into young N2 adult hermaphrodites. After 3-5 days, successfully edited strains were validated using genotyping PCR and sequencing of the edited genomic region.

### RNA-mediated interference (RNAi)

Double stranded RNA (dsRNA) was produced by amplifying DNA templates using primers containing the T7 and T3 promoter sequences (see **Table S2**). Templates were purified using a QIAquick PCR Purification Kit (Qiagen). Single-stranded RNA was generated from DNA templates using MEGAscript T3 and T7 Transcription Kits (Invitrogen) and purified using a MEGAclear Transcription Clean-up Kit (Invitrogen). Single stranded RNA was subsequently annealed at 68°C for 10 minutes followed by 37°C for 30 minutes. L4 hermaphrodites were injected with the final dsRNA products. Injected animals were recovered for 36-48h at 20°C before imaging, except for co-depletion of PP1cβ^GSP-1^ and PP1cα^GSP-2^, and depletion of SDS-22 where the animals were recovered for 24 h prior to imaging. For embryonic lethality assays, L4 stage worms injected with dsRNAs were recovered at 20°C for 24 h, singled onto 35 mm plates, allowed to lay progeny for an additional 24 h before the mothers were transferred to another plate. The next day, the total number of dead embryos and living larvae per plate were scored (24-48h embryo viability). In specific experiments, the mothers were transferred again to another plate, allowed to lay progeny for an additional 24 h prior to scoring (48-72h embryo viability).

### Immunoblotting of C. elegans extracts

Between 50-60 adult worms per condition were transferred to tubes containing M9 buffer (22 mM KH2PO4, 42 mM Na2HPO4, 86 mM NaCl, and 1 mM MgSO4•7H2O), washed 4 times with M9 containing 0.1% Tween-20 and then resuspended in 2X sample buffer (116.7 mM Tris-HCl pH 6.8, 3.3% SDS, 200 mM DTT, 10% glycerol, bromophenol blue). After lysing by sonication followed by boiling, the equivalent of 8-10 worms was loaded onto 4-12% NuPAGE Bis-Tris Gels (Invitrogen). Proteins were then transferred to nitrocellulose membranes, probed with primary antibodies and detected using horseradish (HRP)– conjugated secondary antibodies and WesternBright Sirius (Advansta) chemiluminescent substrate. Membranes were imaged using a ChemiDoc MP imaging system (BioRad). Antibodies used were: goat anti-GFP (gift from D. Drechsel) and mouse anti-actin (Millipore). Secondary antibodies were: HRP-conjugated donkey anti-goat IgG (Jackson Immunoresearch) and HRP-conjugated donkey anti-mouse IgG (Jackson Immunoresearch).

### Fluorescence imaging of C. elegans embryos

Embryos were dissected from adult hermaphrodites into M9 buffer and mounted on a 2% agarose pad. Embryos were then covered with a 22×22 mm coverslip (No. 1.5), and all imaging was performed in a temperature-controlled chamber or a room cooled to 20°C.

Time-lapse imaging of one-cell embryos expressing GFP::H2B and γ-tubulin::GFP was performed on a widefield deconvolution microscope (DeltaVision Elite; Applied Precision, controlled by softWoRx, equipped with a charge-coupled device camera–pco.edge 5.5 sCMOS– and an Olympus 60x 1.42NA PlanApo N/UIS2 objective). A 5 x 2 μm stack was acquired at 10s intervals with 10% illumination intensity (on an InsightSSI illuminator) and 100 ms exposure for GFP.

Imaging of one-cell embryos expressing GFP::PP1cβ^GSP-1^, GFP::PP1cα^GSP-2^ and mStayGold::PP1cα^GSP-2^, as well as mitotic timing analysis of one-cell embryos expressing GFP::H2B and γ-tubulin::GFP were conducted on either an Andor Revolution XD Confocal System (Andor Technology), controlled by Andor iQ3 software, and coupled with a spinning disk confocal scanner unit (CSU-10, Yokogawa) mounted on an inverted microscope, 60x 1.4NA Plan Apochromat lenses, and outfitted with an electron multiplication back-thinned charged coupled device camera (iXon, Andor Technology), or on a Nikon Ti2 confocal system controlled by NIS-Elements software with a Yokogawa CSU-X1 confocal scanner using a 60x 1.42 NA Plan Apochromat Lens and either an Andor iXon electron multiplication back-thinned charge-coupled device (EMCCD) camera or a Kinetix 22 active-pixel sensor (CMOS). 7 x 2 μm or 11 x 2 µm z-stacks were collected every 10 seconds for mitotic timing assays or every 20 seconds for fluorescence localization assays.

To monitor chromosome alignment, one-cell embryos expressing GFP::H2B and γ-tubulin::GFP were imaged on the Nikon Ti2 microscope equipped with a Yokogoawa CSU-X1 spinning disk and an Andor iXon EMCCD camera with a 60x 1.4 NA Plan Aprochromat Lens. 7 x 2 μm image z-stacks were acquired every 2 or 4 seconds.

### Oocyte Meiosis imaging

Oocytes were dissected in meiosis media (25 mM HEPES, 0.5 mg/ml Inulin, 60%Leibovitz L-15 medium, and 20% fetal bovine serum) and mounted on a 2% agarose pad made with egg salt solution (118mM NaCl, 40mM KCl, 3.4mM MgCl_2_, 3.4mM CaCl_2_ and 5mM HEPES (pH 7.4)). Mounted oocytes were imaged with 9 x 1.5um z-stack every 20s on a Nikon Ti2 confocal system with a Yokogawa CSU-X1 confocal scanner using a 100x 1.45NA Plan Apochromat Oil objective lens and a Kinetix 22 active-pixel sensor (CMOS) camera using 12-bit settings. Metaphase (T=0s) was defined as the first frame (20s) after spindle rotation.

### Image analysis

ImageJ (FIJI) was used to process all microscope images. NEBD was scored as the frame where free histone signal in the nucleus equilibrated with the cytoplasm, which was just before abrupt chromosome movements were evident. Anaphase onset was scored as the first frame with visible separation of sister chromatids.

For analysis of chromosome decondensation, nuclear widths were measured by drawing a line across the H2b fluorescence signal of each decondensing chromatin mass at defined time points following anaphase onset and the average of the two values measured per embryo was recorded.

To quantify total GFP::PP1cβ^GSP-1^, GFP::PP1cα^GSP-2^ and mStayGold::PP1cα^GSP-2^signal intensities, an area was drawn surrounding the whole embryo and the mean pixel intensity was measured; background was subtracted by copying the same area to a region without any embryos. To correct for embryo autofluorescence, a strain expressing mCherry-tagged Histone H2b and no green fluorescent protein was imaged under identical exposure conditions as the *in situ*-tagged PP1c strains; the average autofluorescence levels measure for these embryos was subtracted as described in *Fig. S1B*. To quantify cytoplasmic mStayGold::PP1cα^GSP-2^signal in oocytes, an area of a defined size was drawn within the oocyte (ROI) and the mean pixel intensity was measured from three independent ROIs within the oocyte; these three values were averaged and background was subtracted by copying the same area to a region without oocytes. Autofluorescence was not subtracted for the analysis of cytoplasmic fluorescence in meiotic oocytes. Measured values were normalized relative to the mean value of day-matched controls: No RNAi for the *Inh2^SZY-2^(RNAi)* or the WT transgene for RAHF and RAHF-HYNE mutants. To quantify mStayGold::PP1cα^GSP-2^signal on the meiotic half-spindles, mean pixel intensity was measured for the half spindle facing the oocyte cytoplasm at T=0 (metaphase I) and background was subtracted by copying the same area to a cytoplasmic region of the oocyte. To correct for the imaging depth, the mean pixel intensity for mCherry-tagged Histone H2b for each half-spindle was similarly measured and the ratio of mStayGold::PP1cα^GSP-2^ to mCherry-tagged Histone H2b for each half-spindle was calculated. Embryos with very low mCherry::H2b signal (<10 counts mean pixel intensity after background subtraction) were not included in the analysis. Measured ratios for Inh2^SZY-2^ RNAi were normalized relative to the mean ratio of day-matched “No RNAi” controls. For analysis of Inh2^SZY-2^ mutants, calculated mStayGold::PP1cα^GSP-2^ to mCherry-tagged Histone H2b ratios for WT transgene controls were pooled over multiple filming sessions and the mean value employed to normalize the measured ratios for WT, RAHF and RAHF-HYNE transgenes following endogenous Inh2^SZY-2^ depletion.

To monitor chromosome alignment, embryo images were first oriented in the anterior-posterior axis. Chromosome span was measured in movies with 7 x 2 μm z-stacks acquired every 2s or 4s. Stacks were maximum intensity projected, background subtracted, and converted to 8-bit in ImageJ. A minimum bounding box was fit to the edge of the H2b fluorescence signal (the boundary where the pixel value is 0), and the width of the bounding box was measured at 4 second intervals between NEBD and anaphase onset.

### Yeast two-hybrid analysis

Yeast-two hybrid assays were performed by cloning cDNAs encoding GFP::PP1cα^GSP-2^ into the pGBKT7 vector, and Inh2^szy-2^ (either wild-type or SILK or HYNE or RAHF or RAHF-HYNE double mutant) into pGADT7. Diploids with both plasmids were selected on-Leu-Trp medium and tested for interaction on-Leu-Trp-His triple dropout plates.

### AlphaFold3 Structure Predictions

Structure predictions of Inh2^SZY-2^ and the two PP1cs were performed using Alphfold3 (Abramson et al., 2024). PAE plots were generated using the PAEViewer web server (Elfmann and Stulke, 2023). Structural analysis and depictions were carried out in PyMol (DeLano, 2002).

### Statistical analyses

Statistical analysis was performed with Prism 10 software (GraphPad). Analysis of statistical significance was determined using Mann-Whitney tests. Definitions for p-values are provided in the corresponding figure legends. Error bars are 95% confidence intervals.

## ACKNOWLEDGEMENTS

We thank Kevin O’Connell (NIH) for sharing strains and antibodies, and Pablo Lara-Gonzalez (UC Irvine) for helpful discussions. This work was supported by NIH grants to AD (R01GM074215) and KO (R01GM074207).

## AUTHOR CONTRIBUTIONS

The project was conceived by N.V. and A.D. and the majority of experimental effort and analysis was conducted by N.V. A.J.S. conducted the meiosis imaging experiments and J.L.M. conducted the high time-resolution chromosome alignment analysis. The manuscript and figures were drafted by N.V. and A.D. and finalized following feedback from K.O., A.J.S., and J.L.M. A.D. and K.O. supervised the work and obtained funding.

## DATA AVAILABILITY STATEMENT

Data are available in the primary article and in the supplementary materials. Original data, *C. elegans* strains, and plasmids generated in this study are available upon request from the corresponding authors.

## SUPPLEMENTAL MOVIE LEGENDS

**Movie S1: Effects of PP1 catalytic subunit isoform depletions in one-cell *C. elegans* embryos** Timelapse movie of one-cell embryos expressing fluorescently-tagged histone H2b and γ-tubulin, which label chromosomes and spindle poles respectively, for the indicated conditions. 5 x 2 µm Z-stacks were acquired at 10-s intervals and maximum intensity projections generated for each timepoint. Movies are time-aligned with nuclear envelope breakdown (NEBD). Playback rate is 4 frames per second, or 40x relative to real time. Related to Figure 1.

**Movie S2: Localization of the two PP1c isoforms during the first embryonic division** Movies of one-cell embryos expressing *in situ* mStayGold-tagged PP1cα^GSP-2^ (*top*) or GFP-tagged PP1cβ^GSP-1^ (*bottom*). Z-stacks (7 x 2 µm for mSG:: PP1cα^GSP-2^ and 11 x 2 µm for GFP:: PP1cβ^GSP-1^) were acquired at 20-s intervals and maximal intensity projections generated for each timepoint. Playback rate is 4 frames per second, or 80x relative to real time. Related to Figure 1.

**Movie S3: Localization of the two PP1c isoforms during oocyte meiosis I** Timelapse movies of oocytes undergoing meiosis I segregation expressing *in situ* mStayGold(mSG)-tagged PP1cα^GSP-2^ (*top*) or GFP-tagged PP1cβ^GSP-1^ (*bottom*). Both *in situ*-tagged strains expressed mCherry-tagged histone H2b to visualize chromosomes; the mSG::PP1cα^GSP-2^ strain also expressed a mCherry-tagged plasma membrane marker. 9 x 1.5 µm Z-stacks were acquired at 20-s intervals and maximum intensity projections generated for each timepoint. Playback rate is 4 frames per second, or 80x relative to real time. Related to Figure 1.

**Movie S4: Effect of Inh2^SZY-2^ depletion on chromosome alignment and anaphase onset**High time-resolution time-lapse movies monitoring mitotic chromosome dynamics for the indicated conditions. 7 x 2 µm Z-stacks were acquired at 2-s intervals and maximum intensity projections generated for each timepoint. Movies are time-aligned with NEBD. Playback rate is 7 frames per second, or 14x relative to real time. Related to Figure 2 and Figure S2.

**Movie S5: Effect of co-depletion of Inh2^SZY-2^ and PP1c**α**^GSP-2^ on chromosome decondensation** Timelapse movie of one-cell embryos expressing fluorescently-tagged histone H2b and γ-tubulin, which label chromosomes and spindle poles, respectively, for the indicated conditions. Movies are time-aligned with the onset of anaphase and show defects in reforming nuclei. 5 x 2 µm stacks were acquired at 10-s intervals and maximum intensity projections generated for each timepoint. Playback rate is 4 frames per second, or 40x relative to real time. Related to Figure 2.

**Movie S6: Effect of co-depletion of Inh2^SZY-2^ and PP1c**β**^GSP-1^ on chromosome decondensation** Timelapse movie of one-cell embryos expressing fluorescently-tagged histone H2b and γ-tubulin, which label chromosomes and spindle poles, respectively, for the indicated conditions. Movies are time-aligned with the onset of anaphase and show defects in reforming nuclei. 5 x 2 µm stacks were acquired at 10-s intervals and maximum intensity projections generated for each timepoint. Playback rate is 4 frames per second, or 40x relative to real time. Related to Figure 2.

**Movie S7: Consequences of disrupting individual PP1c binding motifs of Inh2^SZY-2^** Movies of one-cell embryo mitosis expressing fluorescently-tagged histone H2b and γ-tubulin for the indicated Inh2^SZY-2^ variants encoded by RNAi-resistant transgenes; endogenous Inh2^SZY-2^ was depleted using RNAi. 7 x 2 µm stacks were acquired at 20-s intervals and maximum intensity projections generated for each timepoint. Movies are time-aligned with respect to NEBD. Playback rate is 4 frames per second, or 80x relative to real time. Related to Figure 3.

**Movie S8: Consequences of disrupting the RVxF, HYNE or both PP1c binding motifs of Inh2^SZY-2^** Movies of one-cell embryo mitosis expressing fluorescently-tagged histone H2b and γ-tubulin for the indicated Inh2^SZY-2^ variants encoded by RNAi-resistant transgenes; endogenous Inh2^SZY-2^ was depleted using RNAi. 7 x 2 µm stacks were acquired at 20-s intervals and maximum intensity projections generated for each timepoint. Movies are time-aligned with respect to NEBD. Playback rate is 4 frames per second, or 80x relative to real time. Related to Figure 4.

---

**Figure S1.**
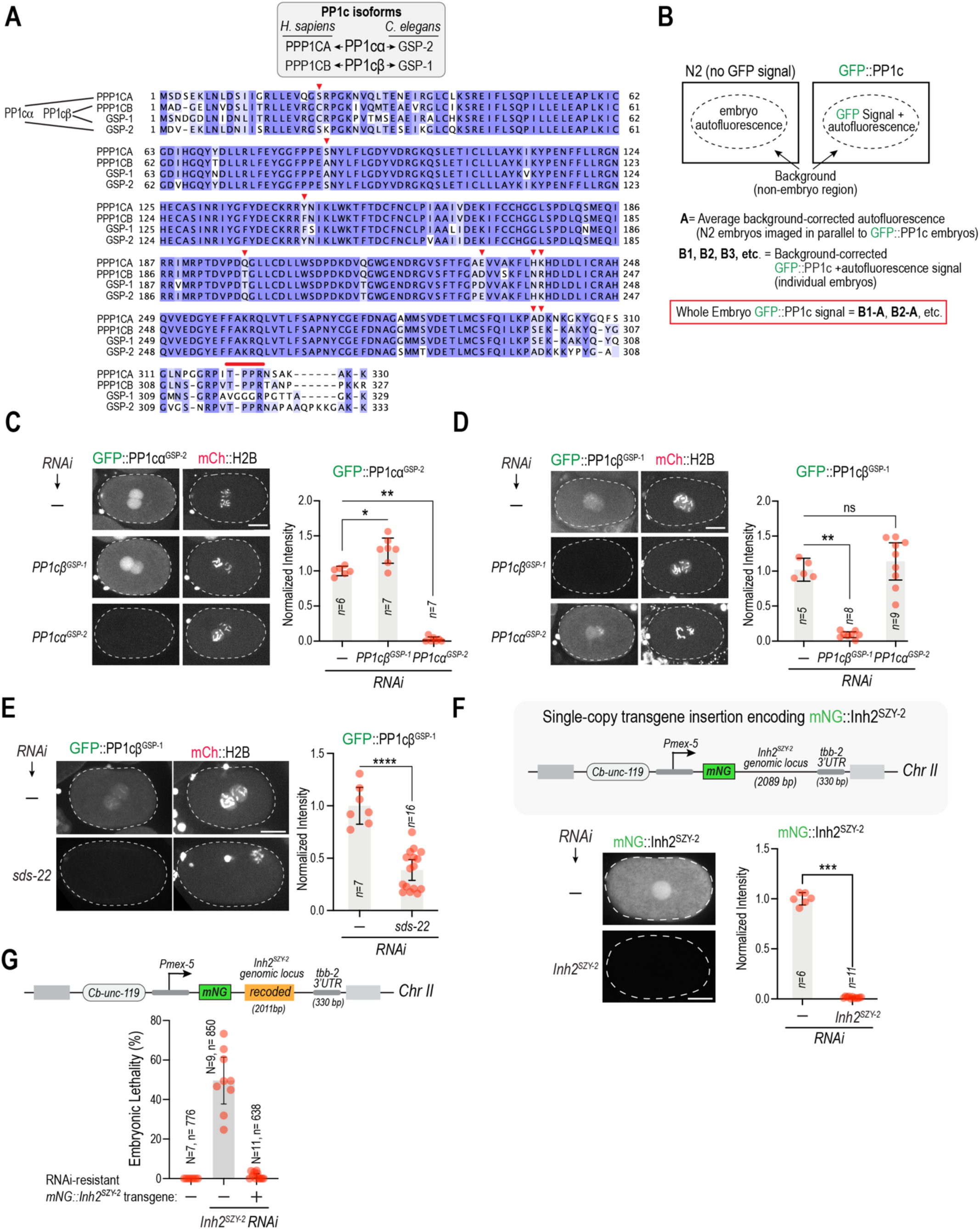
Sequences of two PP1c isoforms expressed in *C. elegans* embryos, evidence of isoform-specific RNAi and efficacy of Inh2^SZY-2^ depletion. **(A)** Sequence alignment of GSP-1 and GSP-2, the two *C. elegans* PP1c catalytic subunits expressed in embryos, with human PPP1CA (PP1α) and PPP1CB (PP1β). Red arrowheads highlight sequence positions supporting GSP-1 similarity to PPP1CB and GSP-2 similarity to PPP1CA. Line highlights the “TPPR” motif that is present in PP1cα^GSP-2^ but not PP1cβ^GSP-1^. **(B)** Schematic describing measurement of whole-embryo GFP fluorescence taking into consideration embryo autofluorescence. The “No GFP” wildtype embryos were imaged in parallel under identical exposure conditions. **(C) & (D)** Images (*left)* and quantification (*right*) of embryo fluorescence, normalized relative to the mean value of the “No *RNAi*” controls, for the indicated conditions. Scale bars, 10 µm. **(E)** Example images and quantification of effect of SDS-22 depletion on PP1cβ^GSP-1^ levels. Note that penetrant SDS-22 depletion results in sterility; thus, the embryos analyzed are likely partially depleted. Scale bar, 10 µm. **(F)** (*top*) Schematic of single copy transgene insertion employed to express mNG-tagged Inh2^SZY-2^. The endogenous *szy-2* coding region is present in this transgene without any alteration. (*bottom*) Evidence of RNAi efficacy, based on quantification of fluorescence of transgene-expressed mNG::Inh2^SZY-2^. Scale bar, 10 µm. *n* in graphs represents number of embryos imaged and analyzed. Error bars are the 95% confidence intervals. p-values are from Mann-Whitney tests (ns=not significant; *=p<0.05; **=p<0.01; ***=p<0.001; ****=p<0.0001). **(G)** Quantification of embryonic lethality for the indicated conditions. For the Inh2^SZY-2^ depletions, lethality of embryos laid 24-48h after injection of dsRNA into late L4 stage larvae was scored. *N* is the number of worms analyzed and *n* the total number of embryos scored.

**Figure S2.**
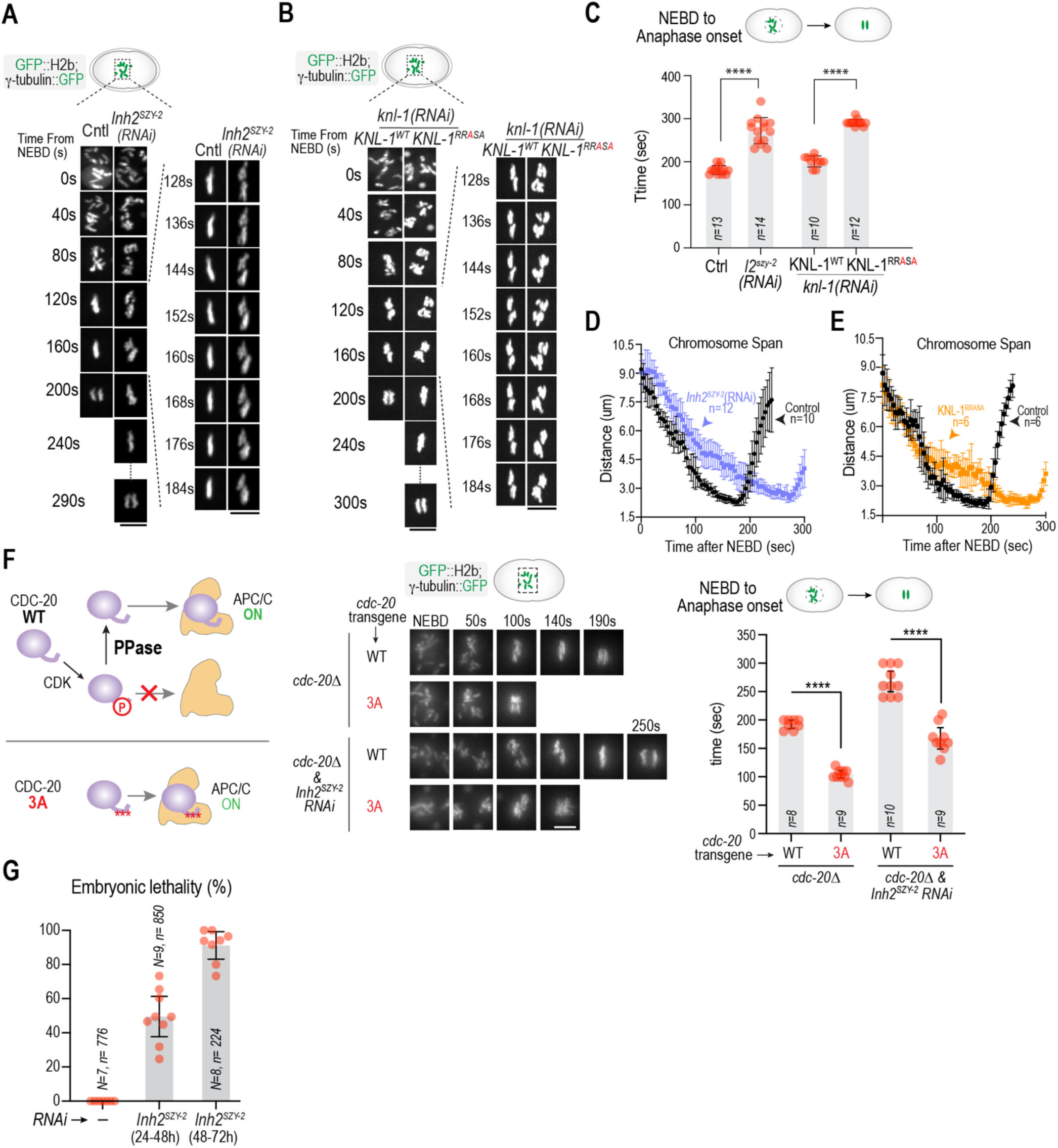
Analysis of chromosome alignment and anaphase onset in Inh2^SZY-2^-depleted embryos. **(A) & (B)** High time-resolution imaging of chromosome dynamics for the indicated conditions. The WT and PP1-docking RRASA mutant versions of KNL-1, encoded by RNAi-resistant transgene insertions, were previously described (Espeut et al., 2012). Both depleting Inh2^SZY-2^- and preventing KNL-1-mediated kinetochore docking of PP1cs results in delayed chromosome alignment and anaphase onset. Scale bars, 5 µm. **(C)** Quantification of the NEBD to anaphase onset interval for the indicated conditions. Note that the Control (no RNAi) and Inh2^SZY-2^ depletion data are the same as in Fig. 2A. *n* is number of embryos analyzed. Error bars are the 95% confidence interval. p-values are from Mann-Whitney tests (****=p<0.0001). **(D) & (E)** Quantification of chromosome span on the spindle, measured as described previously (Cheerambathur et al., 2017) from high-time resolution imaging data of the type shown in panel A. Error bars are the 95% confidence interval. **(F)** (*left*) Schematics of CDC-20 phosphoregulation. (*top*) CDK phosphorylation in its N-terminal tail prevents its binding and activation of the anaphase-promoting complex/cyclosome (APC/C). Phosphatases, including kinetochore-docked PP1c and potentially also cytoplasmic PP1c, dephosphorylate CDC-20, enabling it to bind and activate the APC/C. (*bottom*) A non-phosphorylatable (3A) mutant of CDC-20 is able to bind and activate the APC/C, independently of phosphatase action. (*middle*) Images from timelapse movies for the indicated conditions. The WT and 3A mutant versions of CDC-20 were previously described (Kim et al., 2017). (*right*) Quantification of the NEBD-anaphase onset interval for the indicated conditions. *n* is number of embryos analyzed. p-values are from Mann-Whitney tests (****=p<0.0001). The 3A mutant of CDC-20 significantly reduces the anaphase onset delay caused by Inh2^SZY-2^ depletion. The NEBD-anaphase onset interval of CDC-20 (3A) + Inh2^SZY-2^ depletion is however not identical to CDC-20 (3A), likely because the chromosome alignment defect caused by Inh2^SZY-2^ depletion leads to mild spindle checkpoint activation (the CDC-20 (3A) mutant is competent for spindle checkpoint signaling (Kim et al., 2017). Scale bars, 5 µm. **(G)** Quantification of embryonic lethality for the indicated conditions. For the Inh2^SZY-2^ depletion, lethality of embryos laid 24-48h and 48-72h after injection of dsRNA into late L4 stage larvae was scored separately by moving the injected mother onto new plates at 48h. The later time interval is generally associated with more penetrant depletion of maternally loaded products. *N* is the number of worms analyzed and *n* the total number of embryos scored.

**Figure S3.**
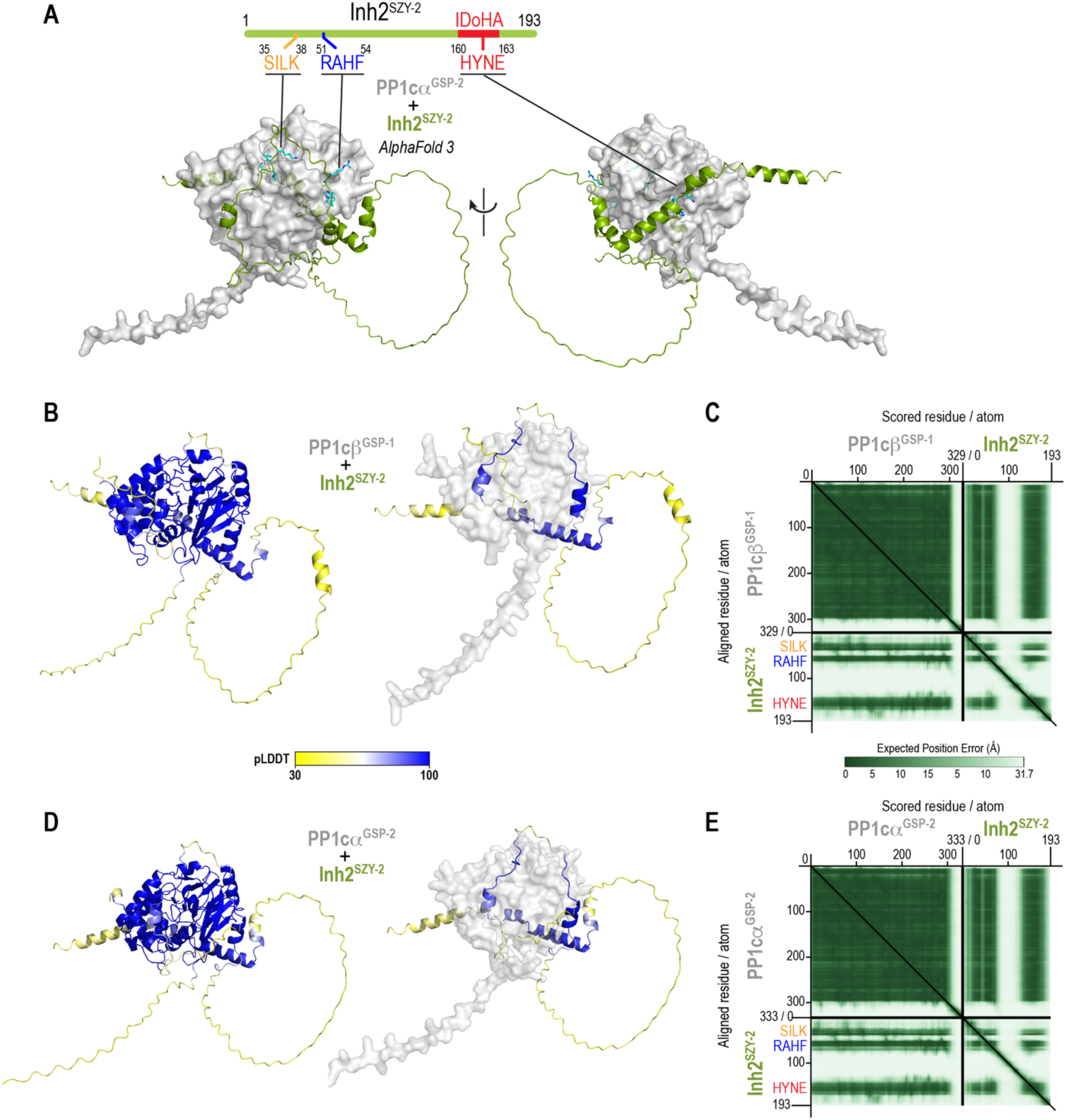
Alphafold 3 models of Inh2^SZY-2^ and *C. elegans* PP1cs. **(A)** Alphafold 3 model of the PP1cα^GSP-2^–Inh2^SZY-2^ complex, highlighting the 3 key PP1c-interacting motifs of Inh2^SZY-2^. **(B) & (D)** (*left*) Models of the PP1c–Inh2^SZY-2^ complexes colored by confidence (pLDDT: predicted local distance difference test). (*right*) Models of the PP1c–Inh2^SZY-2^ complexes where only the Inh2^SZY-2^ chain is colored by confidence (pLDDT: predicted local distance difference test). **(C) & (E)** AlphaFold 3 predicted aligned error (PAE) plots for PP1c–Inh2^SZY-2^ complexes. The plots shown were generated using the PAE Viewer web server.

**Figure S4.**
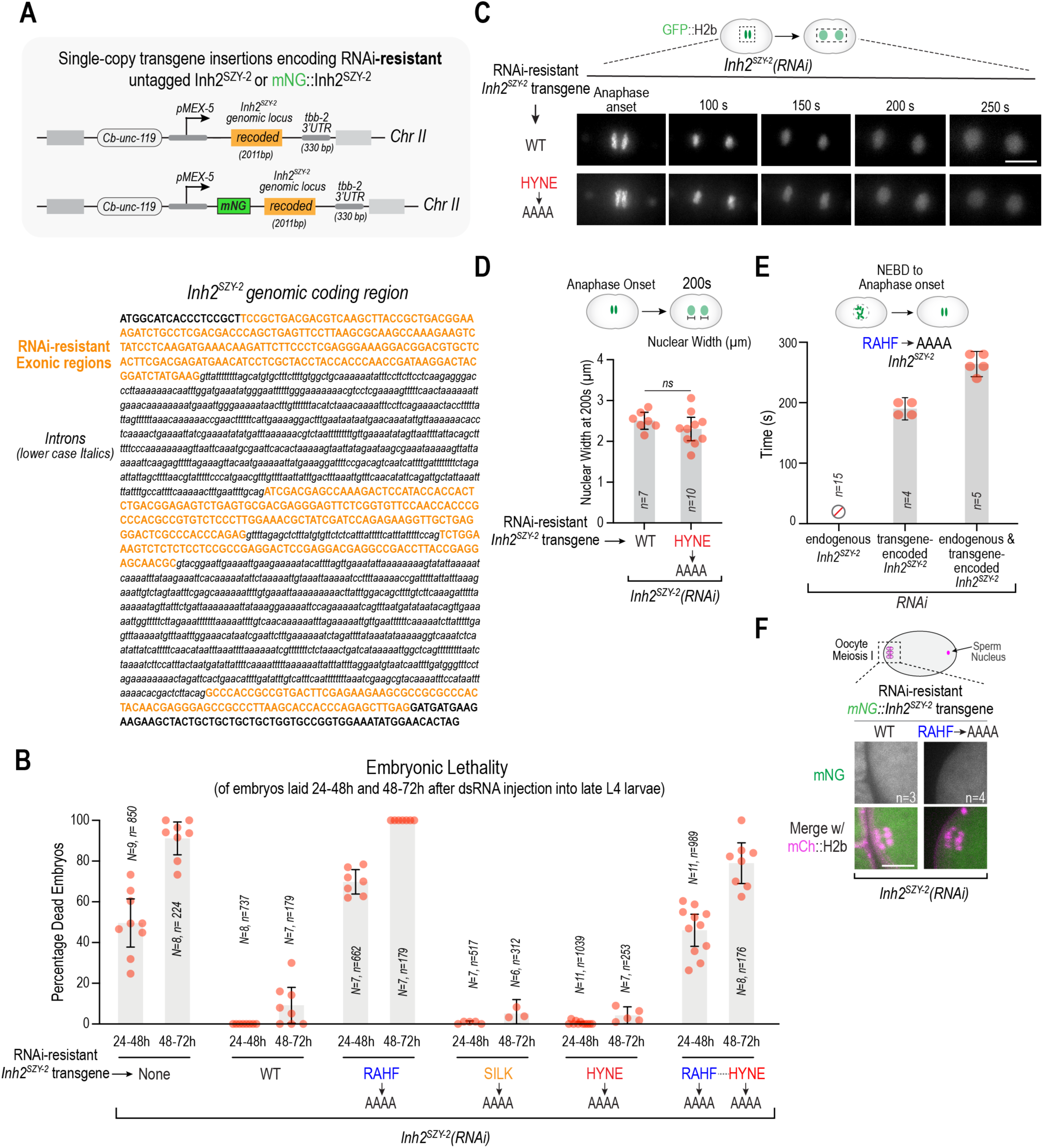
Details on the RNAi-resistant Inh2^SZY-2^ transgene and additional analysis of mutants in PP1c interaction motifs. **(A)** (*top*) Schematic of untagged and mNeonGreen (mNG)-tagged versions of the single copy targeted transgene insertions employed to express WT or various mutant forms of Inh2^SZY-2^. Note that the germline *mex-5* promoter and *tbb-2* 3’ UTRs were used to drive expression. **(B)** Embryonic lethality analysis for the indicated conditions. Lethality of embryos laid 24-48h and 48-72h after injection of dsRNA into late L4 stage larvae was scored separately by moving the injected mothers onto new plates at 48h. The later time interval is generally associated with more penetrant depletion of maternally loaded products. *N* is the number of worms analyzed and *n* the total number of embryos scored. **(C)** Images of chromosome decondensation after anaphase onset for the indicated conditions. Scale bar, 5 µm. **(D)** Quantification of chromosome decondensation by measurement of nuclear width 200s after anaphase onset. Error bars are the 95% confidence interval. p-value is from a Mann-Whitney test (ns = not significant). **(E)** Measurement of NEBD-anaphase onset interval for the indicated RNAi conditions in the strain expressing the RVxF (RAHF->AAAA) mutant Inh2^SZY-2^. When endogenous Inh2^SZY-2^ is depleted, the severity of the phenotype prevents measurement of the NEBD-anaphase onset interval (left-most condition, which is the same as in Fig. 3C). **(F)** Images of the oocyte meiotic spindle region for the indicated conditions. The RVxF mutant form of Inh2^SZY-2^ does not ectopically localize to the spindle region, suggesting it is not forming aberrant trimeric complexes with PP1c and PP1c adapters. The strains imaged also expressed an mCherry-tagged plasma membrane marker. Scale bar, 5 µm.

**Table S1.** List of strains used in this study.

| Strain | Genotype |
| --- | --- |
| N2 | Ancestral |
| OD1702 | *unc-119(ed3)III*; *ItSi560 [oxTi365; oxTi365; pPLG014; Pmex-5::GFP::his-11::tbb-2 3'UTR, tbg-1::gfp::tbb-2 3'UTR; cb-unc-119(+)]V* |
| OD388 | *unc-119(ed3) III*; *ItSi1 [pOD809/pJE110; Pknl-1::KNL-1reencoded::mCherry; cb-unc-119(+)]II*; *ruls32 [pAZ132; pie-1/GFP::his-58; unc-119(+)] III*; *ddls6 [pie-1/GFP::tbg-1; unc-119(+)]V* |
| OD389 | *unc-119(ed3) III*; *ItSi9 [pOD831/pJE120; Pknl-1::KNL-1reencoded(RRASA)::mCherry; cb-unc-119(+)]II*; *ruls32 [pAZ132; pie-1/GFP::his-58; unc-119(+)] III*; *ddls6 [pie-1/GFP::tbg-1; unc-119(+)]V* |
| OD2692 | *ItSi805[pPLG042; Pfzy-1::fzy-1::fzy-1 3'UTR; cb-unc-119(+)]I*; *fzy-1(It20::loxP)II*; *unc-119(ed3)?III*; *ItSi560 [oxTi365; pPLG014; Pmex-5::GFP::his-11::tbb-2 3'UTR, tbg-1::gfp::tbb-2 3'UTR; cb-unc-119(+)]V* |
| OD2918 | *ItSi965[pPLG058; Pfzy-1::fzy-1 T7A, T32A, S87A::fzy-1 3'UTR; cb-unc-119(+)]I*; *fzy-1(It20::loxP)II*; *unc-119(ed3)? III*; *ItSi560 [oxTi365; pPLG014; Pmex-5::GFP::his-11::tbb-2 3'UTR, tbg-1::gfp::tbb-2 3'UTR; cb-unc-119(+)]V* |
| OD3421 | *unc-119(ed3)III?*; *ItSi37 [pAA64; pie-1/mCHERRY::his-58; unc-119 (+)] IV*; *gsp-1(It94[gfp::gsp-1])V* |
| OD2879 | *gsp-2(It27[GFP::gsp-2]) unc-119(ed3) III?*; *ItSi37 [pAA64; pie-1/mCHERRY::his-58; unc-119 (+)] IV* |
| OD5844 | *ItSi1972 [pNV48/pOD5544; Pmex-5::szy-2 Reencoded::tbb-2 3'UTR; cb-unc-119(+)]II*; *unc-119(ed3)III* |
| OD5745 | *ItSi1942[pNV23/pOD5540; Pmex-5::szy-2 Reencoded::RAHF>AAAA::tbb-2 3'UTR; cb-unc-119(+)]II*; *unc-119(ed3)III* |
| OD5751 | *ItSi1948[pNV33/pOD5542; Pmex-5::szy-2 Reencoded::SILK>AAAA::tbb-2 3'UTR; cb-unc-119(+)]II*; *unc-119(ed3)III* |
| OD5943 | *ItSi2000 [pNV57/ pOD5546; Pmex-5::szy-2 Reencoded::RAHF>AAAA_HYNE>AAAA::tbb-2 3'UTR; cb-unc-119(+)]II* ; *unc-119(ed3)III* |
| OD5775 | *ItSi1953[pNV35/pOD5543; Pmex-5::szy-2 Reencoded::HYNE>AAAA::tbb-2 3'UTR; cb-unc-119(+)]II*; *unc-119(ed3)III* |
| OD6062 | *gsp-2(It266[mStayGold::gsp-2]) III* |
| OD6124 | *ItSi1088[oxTi185; pMO005; Pmex-5::mCherry::ph::tbb-2 3'-UTR::operon linker::mCherry::his-11::tbb-2 3'-UTR]I*; *gsp-2(It266[mStayGold::gsp-2]) III*; *unc-119(ed3)III ?* |
| OD5565 | *ItSi1928[pNV12/pOD5537; Pmex-5::mNG::szy-2::tbb-2 3'UTR; cb-unc-119(+)]II*; *unc-119(ed3)III* |
| OD5566 | *ItSi1929[pNV14/pOD5538; Pmex-5::mNeonGreen::szy-2 Reencoded::tbb-2 3'UTR; cb-unc-119(+)]II*; *unc-119(ed3)III* |
| OD5839 | *ItSi1929[pNV14/pOD5538; Pmex-5::mNeonGreen::szy-2 Reencoded::tbb-2 3'UTR; cb-unc-119(+)]II*; *unc-119(ed3)III?*; *knl-1( It75[knl-1::mCherry]) III* |
| OD5862 | *ItSi1971[pNV48/ pOD5544; Pmex-5::szy-2 Reencoded::tbb-2 3'UTR; cb-unc-119(+)]II*; *unc-119(ed3)III?*; *ItSi560 [oxTi365; oxTi365; pPLG014; Pmex-5::GFP::his-11::tbb-2 3'UTR, tbg-1::gfp::tbb-2 3'UTR; cb-unc-119(+)]V* |
| OD5863 | *ItSi1948[pNV33/ pOD5542; Pmex-5::szy-2 Reencoded::SILK>AAAA::tbb-2 3'UTR; cb-unc-119(+)]II*; *unc-119(ed3)III?*; *ItSi560 [oxTi365; oxTi365; pPLG014; Pmex-5::GFP::his-11::tbb-2 3'UTR, tbg-1::gfp::tbb-2 3'UTR; cb-unc-119(+)]V* |
| OD5865 | *ItSi1953[pNV35/ pOD5543; Pmex-5::szy-2 Reencoded::HYNE>AAAA::tbb-2 3'UTR; cb-unc-119(+)]II*; *unc-119(ed3)III?*; *ItSi560 [oxTi365; oxTi365; pPLG014; Pmex-5::GFP::his-11::tbb-2 3'UTR, tbg-1::gfp::tbb-2 3'UTR; cb-unc-119(+)]V* |
| OD5867 | *ItSi1942[pNV23/ pOD5540; Pmex-5::szy-2 Reencoded::RAHF>AAAA::tbb-2 3'UTR; cb-unc-119(+)]II; unc-119(ed3)III?; ItSi560 [oxTi365; oxTi365; pPLG014; Pmex-5::GFP::his-11::tbb-2 3'UTR, tbg-1::gfp::tbb-2 3'UTR; cb-unc-119(+)]V* |
| OD6207 | *ItSi1088[oxTi185; pMO005; Pmex-5::mCherry::ph::tbb-2 3'-UTR::operon linker::mCherry::his-11::tbb-2 3'-UTR]I; ItSi1940[pNV22/ pOD5539; Pmex-5::mNeonGreen::szy-2 Reencoded::RAHF>AAAA::tbb-2 3'UTR; cb-unc-119(+)]II; unc-119(ed3)III?* |
| OD6208 | *ItSi1088[oxTi185; pMO005; Pmex-5::mCherry::ph::tbb-2 3'-UTR::operon linker::mCherry::his-11::tbb-2 3'-UTR]I; ItSi1929[pNV14/ pOD5538; Pmex-5::mNeonGreen::szy-2 Reencoded::tbb-2 3'UTR; cb-unc-119(+)]II; unc-119(ed3)III?* |
| OD6211 | *ItSi1088[oxTi185; pMO005; Pmex-5::mCherry::ph::tbb-2 3'-UTR::operon linker::mCherry::his-11::tbb-2 3'-UTR]I; ItSi1971[pNV48/ pOD5544; Pmex-5::szy-2 Reencoded::tbb-2 3'UTR; cb-unc-119(+)]II; gsp-2(It266[mStayGold::gsp-2]) III; unc-119(ed3)III ?* |
| OD6212 | *ItSi1088[oxTi185; pMO005; Pmex-5::mCherry::ph::tbb-2 3'-UTR::operon linker::mCherry::his-11::tbb-2 3'-UTR]I; ItSi1942[pNV23/ pOD5540; Pmex-5::szy-2 Reencoded::RAHF>AAAA::tbb-2 3'UTR; cb-unc-119(+)]II; gsp-2(It266[mStayGold::gsp-2]) III; unc-119(ed3)III ?* |
| OD6213 | *ItSi1088[oxTi185; pMO005; Pmex-5::mCherry::ph::tbb-2 3'-UTR::operon linker::mCherry::his-11::tbb-2 3'-UTR]I; ItSi2000 [pNV57/ pOD5546; Pmex-5::szy-2 Reencoded::RAHF>AAAA_HYNE>AAAA ::tbb-2 3'UTR; cb-unc-119(+)]II ; gsp-2(It266[mStayGold::gsp-2]) III; unc-119(ed3)III ?* |
| OD6249 | *ItSi2002 [pNV57/ pOD5546; Pmex-5::szy-2 Reencoded::RAHF>AAAA_HYNE>AAAA ::tbb-2 3'UTR; cb-unc-119(+)]II ; unc-119(ed3)III?; ItSi560 [oxTi365; oxTi365; pPLG014; Pmex-5::GFP::his-11::tbb-2_3'UTR, tbg-1::gfp::tbb-2_3'UTR; cb-unc-119(+)]V* |

**Table S2.**
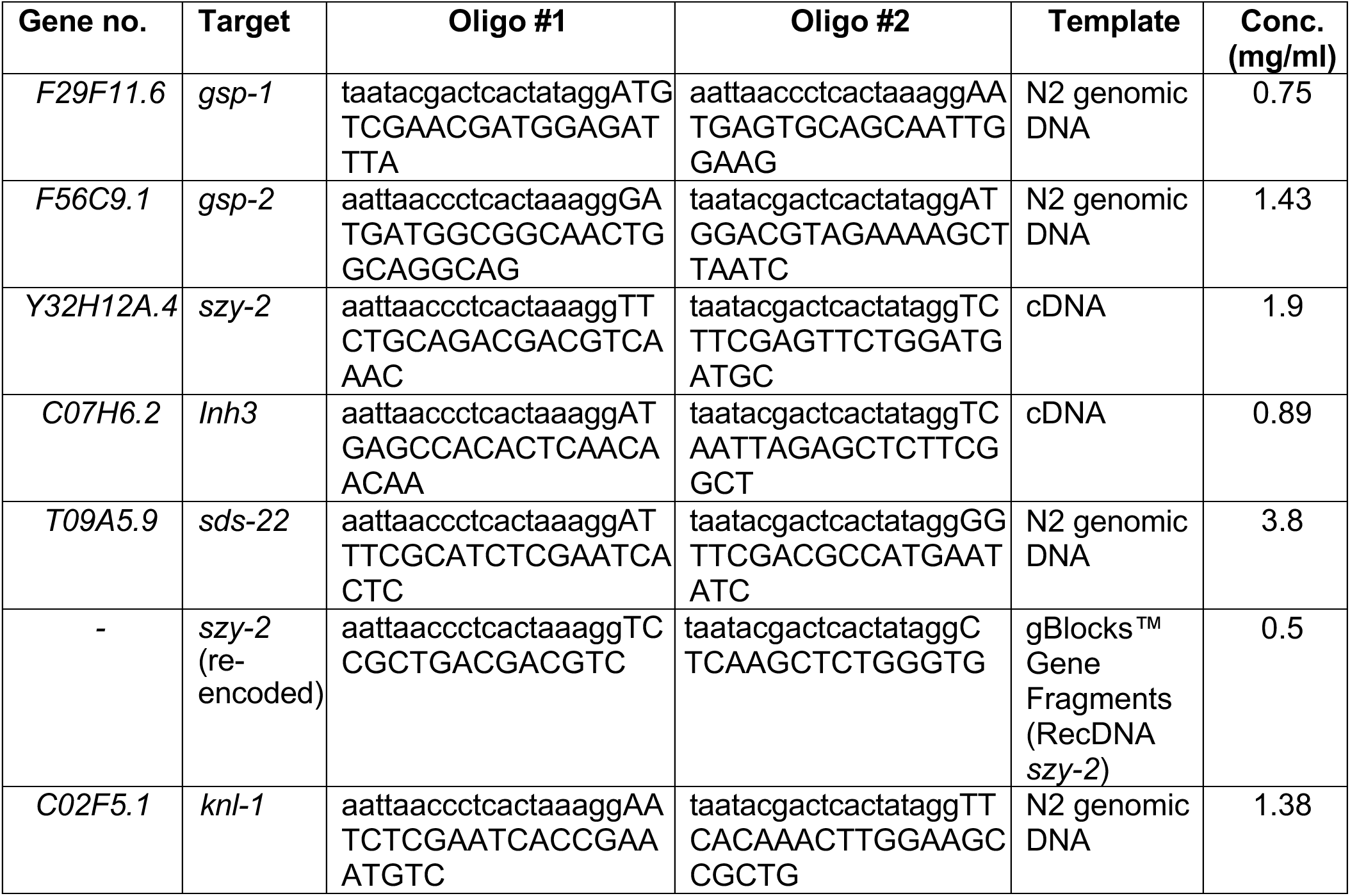
Oligonucleotides used for dsRNA production.

## REFERENCES

Abramson, J., J. Adler, J. Dunger, R. Evans, T. Green, A. Pritzel, O. Ronneberger, L. Willmore, A.J. Ballard, J. Bambrick, S.W. Bodenstein, D.A. Evans, C.C. Hung, M. O’Neill, D. Reiman, K. Tunyasuvunakool, Z. Wu, A. Zemgulyte, E. Arvaniti, C. Beattie, O. Bertolli, A. Bridgland, A. Cherepanov, M. Congreve, A.I. Cowen-Rivers, A. Cowie, M. Figurnov, F.B. Fuchs, H. Gladman, R. Jain, Y.A. Khan, C.M.R. Low, K. Perlin, A. Potapenko, P. Savy, S. Singh, A. Stecula, A. Thillaisundaram, C. Tong, S. Yakneen, E.D. Zhong, M. Zielinski, A. Zidek, V. Bapst, P. Kohli, M. Jaderberg, D. Hassabis, and J.M. Jumper. 2024. Accurate structure prediction of biomolecular interactions with AlphaFold 3. Nature. 630:493–500.

Alessi, D.R., A.J. Street, P. Cohen, and P.T. Cohen. 1993. Inhibitor-2 functions like a chaperone to fold three expressed isoforms of mammalian protein phosphatase-1 into a conformation with the specificity and regulatory properties of the native enzyme. Eur J Biochem. 213:1055–1066.

Antonin, W., and H. Neumann. 2016. Chromosome condensation and decondensation during mitosis. Curr Opin Cell Biol. 40:15–22.

Ballou, L.M., D.L. Brautigan, and E.H. Fischer. 1983. Subunit structure and activation of inactive phosphorylase phosphatase. Biochemistry. 22:3393–3399.

Bollen, M. 2001. Combinatorial control of protein phosphatase-1. Trends Biochem Sci. 26:426–431.

Bollen, M., W. Peti, M.J. Ragusa, and M. Beullens. 2010. The extended PP1 toolkit: designed to create specificity. Trends Biochem Sci. 35:450–458.

Calvi, I., F. Schwager, and M. Gotta. 2022. PP1 phosphatases control PAR-2 localization and polarity establishment in C. elegans embryos. J Cell Biol. 221.

Cao, X., M. Lake, G. Van der Hoeven, Z. Claes, J. Del Pino Garcia, S. Lemaire, E.C. Greiner, S. Karamanou, A. Van Eynde, A.N. Kettenbach, D. Natera de Benito, L. Carrera Garcia, C. Hernando Davalillo, C. Ortez, A. Nascimento, R. Urreizti, and M. Bollen. 2024. SDS22 coordinates the assembly of holoenzymes from nascent protein phosphatase-1. Nat Commun. 15:5359.

Cao, X., S. Lemaire, and M. Bollen. 2022. Protein phosphatase 1: life-course regulation by SDS22 and Inhibitor-3. FEBS J. 289:3072–3085.

Ceulemans, H., and M. Bollen. 2004. Functional diversity of protein phosphatase-1, a cellular economizer and reset button. Physiol Rev. 84:1–39.

Cheerambathur, D.K., B. Prevo, N. Hattersley, L. Lewellyn, K.D. Corbett, K. Oegema, and A. Desai. 2017. Dephosphorylation of the Ndc80 Tail Stabilizes Kinetochore-Microtubule Attachments via the Ska Complex. Dev Cell. 41:424–437 e424.

Choy, M.S., T.M. Moon, R. Ravindran, J.A. Bray, L.C. Robinson, T.L. Archuleta, W. Shi, W. Peti, K. Tatchell, and R. Page. 2019. SDS22 selectively recognizes and traps metal-deficient inactive PP1. Proc Natl Acad Sci U S A. 116:20472–20481.

Cohen, P. 1989. The structure and regulation of protein phosphatases. Annu Rev Biochem. 58:453–508.

Cohen, P.T. 2002. Protein phosphatase 1--targeted in many directions. J Cell Sci. 115:241–256.

Dancheck, B., M.J. Ragusa, M. Allaire, A.C. Nairn, R. Page, and W. Peti. 2011. Molecular investigations of the structure and function of the protein phosphatase 1-spinophilin-inhibitor 2 heterotrimeric complex. Biochemistry. 50:1238–1246.

DeLano, W.L. 2002. PyMOL: An Open-Source Molecular Graphics Tool. CCP4 Newsletter on ProteinCrystallography 44–53.

Dohadwala, M., E.F. da Cruz e Silva, F.L. Hall, R.T. Williams, D.A. Carbonaro-Hall, A.C. Nairn, P. Greengard, and N. Berndt. 1994. Phosphorylation and inactivation of protein phosphatase 1 by cyclin-dependent kinases. Proc Natl Acad Sci U S A. 91:6408–6412.

Elfmann, C., and J. Stulke. 2023. PAE viewer: a webserver for the interactive visualization of the predicted aligned error for multimer structure predictions and crosslinks. Nucleic Acids Res. 51:W404–W410.

Espeut, J., D.K. Cheerambathur, L. Krenning, K. Oegema, and A. Desai. 2012. Microtubule binding by KNL-1 contributes to spindle checkpoint silencing at the kinetochore. J Cell Biol. 196:469–482.

Eto, M., E. Elliott, T.D. Prickett, and D.L. Brautigan. 2002. Inhibitor-2 regulates protein phosphatase-1 complexed with NimA-related kinase to induce centrosome separation. J Biol Chem. 277:44013–44020.

Freville, A., K. Cailliau-Maggio, C. Pierrot, G. Tellier, H. Kalamou, S. Lafitte, A. Martoriati, R.J. Pierce, J.F. Bodart, and J. Khalife. 2013. Plasmodium falciparum encodes a conserved active inhibitor-2 for Protein Phosphatase type 1: perspectives for novel anti-plasmodial therapy. BMC Biol. 11:80.

Frokjaer-Jensen, C., M.W. Davis, C.E. Hopkins, B.J. Newman, J.M. Thummel, S.P. Olesen, M. Grunnet, and E.M. Jorgensen. 2008. Single-copy insertion of transgenes in Caenorhabditis elegans. Nat Genet. 40:1375–1383.

Hashimshony, T., M. Feder, M. Levin, B.K. Hall, and I. Yanai. 2015. Spatiotemporal transcriptomics reveals the evolutionary history of the endoderm germ layer. Nature. 519:219–222.

Hattersley, N., D. Cheerambathur, M. Moyle, M. Stefanutti, A. Richardson, K.Y. Lee, J. Dumont, K. Oegema, and A. Desai. 2016. A Nucleoporin Docks Protein Phosphatase 1 to Direct Meiotic Chromosome Segregation and Nuclear Assembly. Dev Cell. 38:463–477.

Hemmings, B.A., T.J. Resink, and P. Cohen. 1982. Reconstitution of a Mg-ATP-dependent protein phosphatase and its activation through a phosphorylation mechanism. FEBS Lett. 150:319–324.

Hendrickx, A., M. Beullens, H. Ceulemans, T. Den Abt, A. Van Eynde, E. Nicolaescu, B. Lesage, and M. Bollen. 2009. Docking motif-guided mapping of the interactome of protein phosphatase-1. Chem Biol. 16:365–371.

Heroes, E., B. Lesage, J. Gornemann, M. Beullens, L. Van Meervelt, and M. Bollen. 2013. The PP1 binding code: a molecular-lego strategy that governs specificity. FEBS J. 280:584–595.

Hirano, M., R. Ando, S. Shimozono, M. Sugiyama, N. Takeda, H. Kurokawa, R. Deguchi, K. Endo, K. Haga, R. Takai-Todaka, S. Inaura, Y. Matsumura, H. Hama, Y. Okada, T. Fujiwara, T. Morimoto, K. Katayama, and A. Miyawaki. 2022. A highly photostable and bright green fluorescent protein. Nat Biotechnol. 40:1132–1142.

Hsu, J.Y., Z.W. Sun, X. Li, M. Reuben, K. Tatchell, D.K. Bishop, J.M. Grushcow, C.J. Brame, J.A. Caldwell, D.F. Hunt, R. Lin, M.M. Smith, and C.D. Allis. 2000. Mitotic phosphorylation of histone H3 is governed by Ipl1/aurora kinase and Glc7/PP1 phosphatase in budding yeast and nematodes. Cell. 102:279–291.

Huang, F.L., and W.H. Glinsmann. 1976. Separation and characterization of two phosphorylase phosphatase inhibitors from rabbit skeletal muscle. Eur J Biochem. 70:419–426.

Hurley, T.D., J. Yang, L. Zhang, K.D. Goodwin, Q. Zou, M. Cortese, A.K. Dunker, and A.A. DePaoli-Roach. 2007. Structural basis for regulation of protein phosphatase 1 by inhibitor-2. J Biol Chem. 282:28874–28883.

Ivorra-Molla, E., D. Akhuli, M.B.L. McAndrew, W. Scott, L. Kumar, S. Palani, M. Mishima, A. Crow, and M.K. Balasubramanian. 2024. A monomeric StayGold fluorescent protein. Nat Biotechnol. 42:1368–1371.

Kim, T., P. Lara-Gonzalez, B. Prevo, F. Meitinger, D.K. Cheerambathur, K. Oegema, and A. Desai. 2017. Kinetochores accelerate or delay APC/C activation by directing Cdc20 to opposing fates. Genes Dev. 31:1089–1094.

Kwon, Y.G., S.Y. Lee, Y. Choi, P. Greengard, and A.C. Nairn. 1997. Cell cycle-dependent phosphorylation of mammalian protein phosphatase 1 by cdc2 kinase. Proc Natl Acad Sci U S A. 94:2168–2173.

Lemaire, S., and M. Bollen. 2020. Protein phosphatase-1: dual activity regulation by Inhibitor-2. Biochem Soc Trans. 48:2229–2240.

Lemaire, S., M. Ferreira, Z. Claes, R. Derua, M. Lake, G. Van der Hoeven, F. Withof, X. Cao, E.C. Greiner, A.N. Kettenbach, A. Van Eynde, and M. Bollen. 2024. PPP1R2 stimulates protein phosphatase-1 through stabilisation of dynamic subunit interactions. Nat Commun. 15:9822.

Li, Y., I. Calvi, and M. Gotta. 2025. SDS-22 stabilizes GSP-1/-2 PP1 subunits contributing to polarity establishment in C. elegans embryos. EMBO Rep. 26:6240–6265.

Moura, M., J. Barbosa, I. Pinto, N. Leca, S. Cunha-Silva, A.E. Verza, P.D. Pedroso, S. Lemaire, J. Duro, M. Simoes-da-Silva, A. Oliveira, R. Reis, J. Nilsson, C. Sunkel, A. Musacchio, M. Bollen, and C. Conde. 2025. Polo kinase inhibits protein phosphatase 1 to promote the spindle assembly checkpoint and prevent aneuploidy. Curr Biol. 35:5289–5307 e5288.

Nadarajan, S., E. Altendorfer, T.T. Saito, M. Martinez-Garcia, and M.P. Colaiacovo. 2021. HIM-17 regulates the position of recombination events and GSP-1/2 localization to establish short arm identity on bivalents in meiosis. Proc Natl Acad Sci U S A. 118.

Nilsson, J. 2019. Protein phosphatases in the regulation of mitosis. J Cell Biol. 218:395–409.

Paix, A., A. Folkmann, D. Rasoloson, and G. Seydoux. 2015. High Efficiency, Homology-Directed Genome Editing in Caenorhabditis elegans Using CRISPR-Cas9 Ribonucleoprotein Complexes. Genetics. 201:47–54.

Park, I.K., and A.A. DePaoli-Roach. 1994. Domains of phosphatase inhibitor-2 involved in the control of the ATP-Mg-dependent protein phosphatase. J Biol Chem. 269:28919–28928.

Peel, N., J. Iyer, A. Naik, M.P. Dougherty, M. Decker, and K.F. O’Connell. 2017. Protein Phosphatase 1 Down Regulates ZYG-1 Levels to Limit Centriole Duplication. PLoS Genet. 13:e1006543.

Peti, W., A.C. Nairn, and R. Page. 2013. Structural basis for protein phosphatase 1 regulation and specificity. FEBS J. 280:596–611.

Qian, J., B. Lesage, M. Beullens, A. Van Eynde, and M. Bollen. 2011. PP1/Repo-man dephosphorylates mitotic histone H3 at T3 and regulates chromosomal aurora B targeting. Curr Biol. 21:766–773.

Trinkle-Mulcahy, L., J. Andersen, Y.W. Lam, G. Moorhead, M. Mann, and A.I. Lamond. 2006. Repo-Man recruits PP1 gamma to chromatin and is essential for cell viability. J Cell Biol. 172:679–692.

Tung, H.Y., W. Wang, and C.S. Chan. 1995. Regulation of chromosome segregation by Glc8p, a structural homolog of mammalian inhibitor 2 that functions as both an activator and an inhibitor of yeast protein phosphatase 1. Mol Cell Biol. 15:6064–6074.

Tzur, Y.B., C. Egydio de Carvalho, S. Nadarajan, I. Van Bostelen, Y. Gu, D.S. Chu, I.M. Cheeseman, and M.P. Colaiacovo. 2012. LAB-1 targets PP1 and restricts Aurora B kinase upon entrance into meiosis to promote sister chromatid cohesion. PLoS Biol. 10:e1001378.

Vagnarelli, P., D.F. Hudson, S.A. Ribeiro, L. Trinkle-Mulcahy, J.M. Spence, F. Lai, C.J. Farr, A.I. Lamond, and W.C. Earnshaw. 2006. Condensin and Repo-Man-PP1 co-operate in the regulation of chromosome architecture during mitosis. Nat Cell Biol. 8:1133–1142.

Vagnarelli, P., S. Ribeiro, L. Sennels, L. Sanchez-Pulido, F. de Lima Alves, T. Verheyen, D.A. Kelly, C.P. Ponting, J. Rappsilber, and W.C. Earnshaw. 2011. Repo-Man coordinates chromosomal reorganization with nuclear envelope reassembly during mitotic exit. Dev Cell. 21:328–342.

van den Boom, J., G. Marini, H. Meyer, and H.R. Saibil. 2023. Structural basis of ubiquitin-independent PP1 complex disassembly by p97. EMBO J. 42:e113110.

Verbinnen, I., M. Ferreira, and M. Bollen. 2017. Biogenesis and activity regulation of protein phosphatase 1. Biochem Soc Trans. 45:89–99.

Virshup, D.M., and S. Shenolikar. 2009. From promiscuity to precision: protein phosphatases get a makeover. Mol Cell. 33:537–545.

Wang, W., C. Cronmiller, and D.L. Brautigan. 2008a. Maternal phosphatase inhibitor-2 is required for proper chromosome segregation and mitotic synchrony during Drosophila embryogenesis. Genetics. 179:1823–1833.

Wang, W., P.T. Stukenberg, and D.L. Brautigan. 2008b. Phosphatase inhibitor-2 balances protein phosphatase 1 and aurora B kinase for chromosome segregation and cytokinesis in human retinal epithelial cells. Mol Biol Cell. 19:4852–4862.

Weith, M., J. Seiler, J. van den Boom, M. Kracht, J. Hulsmann, I. Primorac, J. Del Pino Garcia, F. Kaschani, M. Kaiser, A. Musacchio, M. Bollen, and H. Meyer. 2018. Ubiquitin-Independent Disassembly by a p97 AAA-ATPase Complex Drives PP1 Holoenzyme Formation. Mol Cell. 72:766–777 e766.

Wu, J.C., A.C. Go, M. Samson, T. Cintra, S. Mirsoian, T.F. Wu, M.M. Jow, E.J. Routman, and D.S. Chu. 2012. Sperm development and motility are regulated by PP1 phosphatases in Caenorhabditis elegans. Genetics. 190:143–157.

